# Direct and indirect benefits of cooperation in collective defense against predation

**DOI:** 10.1101/2025.08.19.670848

**Authors:** Raphael Ritter, Heikki Helanterä, Riikka Tynkkynen, Saskia Wutke, Carita Lindstedt

## Abstract

The evolution and maintenance of public goods cooperation, despite cheating, remains a key interest in social biology. Identifying how ecological factors determine the direct and indirect benefits that maintain cooperation has proven challenging, as these can vary significantly across species and environments. Here, we study this problem using the social pine sawfly *Neodiprion sertifer* (Hymenoptera) as a model system. During their larval stage, *N. sertifer* live in groups and collectively secrete a defensive fluid against predators. This behavior comprises a public good as it is costly to exhibit and beneficial to others, and individuals vary in their contribution to group defense. We experimentally manipulated individual contributions to defense to assess how these influence survival under natural insect predation. Our results indicate that defense has a group-level benefit as individuals were more likely to survive in cooperative groups with a higher proportion of defending larvae. Moreover, being able to deploy defensive fluid confers direct survival benefits. Genetic and phenotypic analyses of natural populations further show that kin selection promotes collective defense, as groups of larvae are often composed of full siblings. We also find that the contribution to defense is female-biased and diminishes in larger, more male-biased groups, and to some extent with decreased kinship, indicating that individuals adjust their contributions based on social context. Overall, we find that contribution to the collective defense provides both direct and indirect benefits and that individuals regulate their contributions mainly based on the social environment, resulting in variation within and among natural populations.

**Significance statement:** Individuals in groups can cooperate to achieve shared goals, but they face an evolutionary challenge: if the benefits of cooperation are shared equally, freeloaders receive the same benefits as others while contributing less. Although theoretical solutions to this problem are abundant, we still lack empirical evidence on how those mechanisms function in natural systems. We study this with pine sawfly larvae, which defend collectively against predators. We show that while cooperation increases the survival of individuals and their relatives, they adjust their contributions based on who and how many they are surrounded by, relying more on freeloading in larger and male-biased groups. This results in variation in cooperativeness but prevents freeloaders from taking over.

## Introduction

Individuals can cooperate in public goods which are often costly to produce for an individual, but beneficial for all members of a local group or population (1). Public goods systems are abundant across species with examples including social immunity in burying beetles (2), parasitoid wasps with cooperative brood care (3), siderophores and biofilms in bacteria (4, 5) and communal nesting in birds (6). The fundamental challenge of public goods systems is their vulnerability to exploitation or cheating, i.e., benefiting from the public good without contributing to the production of it (1, 7–9). If contributing is costly, and benefits are shared, it is not a priori guaranteed that contributing to collective good leads to higher fitness than reaping the benefit without having contributed to its production (10, 11). This raises the question of the evolutionary stability of such systems over time.

Several mechanisms that select for cooperation have been identified (1, 11–15), affecting both direct and indirect fitness components of inclusive fitness (16, 17). For example, costs and benefits of exploitation can be limited through negative frequency-dependence (11, 18, 19). In this case, the fitness of cheats will decrease when their frequency increases limiting their spread. The frequency-dependence can become stronger when at least some group members are related (20). In this context, an increase in cheating within a group would increasingly harm the relatives as well. Finally, if more cooperative individuals have better access to the public goods they have contributed to (i.e., direct benefits of cooperation), it should limit the expression of cheating (21–24). While significant progress has been made using microbial systems to study these mechanisms empirically (10, 18, 25–28), these predictions still need to be experimentally validated in more complex study systems where cooperation is facultative, and which show a higher degree of plasticity. This will help us to better determine the sources of diversity in cooperation and its maintenance in nature.

Here, we investigate solutions to the public goods dilemma within a chemical defense framework (29). Many organisms defend against predators using defensive secretions, which are costly to produce and maintain (30–36). Their expression is often facultative (29–31, 36), giving individuals the option to choose whether or not to deploy the defensive fluid when approached by a predator. In general, deploying the fluid may deter the predator, or harm it, limiting its future attacks (29, 37). Furthermore, chemically defended prey are also considered to contribute to public goods by educating predators to avoid prey with similar appearance in future encounters (38, 29, 39). These shared costs of predator education are then a collective production of benefits shared by local group members or the population, i.e., cooperation (29, 40, 41). They apply both to solitary and gregarious species and can also occur at an interspecific level in Müllerian mimicry, i.e., defended individuals belonging to different species share the same warning signal (42).

Group living by the prey has been shown to enhance avoidance learning by predators (43, 44). This is because, when attacking an individual that is part of a group, a predator encounters the chemically defended prey more frequently, making it easier for the predator to recognize and memorize the appearance of that prey type and avoid attacking such prey in future encounters (44–46). A higher potency of chemical defense (47), and a higher proportion of individuals participating in the defense within a group (48, 49) are also expected to increase defense success. Finally, when multiple individuals release secretions simultaneously, the magnitude of these stimuli is likely to increase protection (44, 50, 51), for instance by forming a barrier of defensive fluid that makes it difficult for a predator to reach a single prey without being exposed to the harmful effects of the secretion. In addition to these public goods-type mechanisms, displaying defensive secretions can reduce the risks associated with predation for an individual directly (i.e., providing individual benefits). This is because a predator may assess the unpalatability or toxicity of an individual prey using visual, olfactory or gustatory cues of the defensive secretion, without actually killing the prey (52). Being able to secrete the defensive fluid will also help an individual in “fighting” against the attacking predator.

To study empirically the mechanisms that maintain public goods cooperation in chemical defense, we used social haplodiploid European pine sawflies (*Neodiprion sertifer*) as a model system. They live solitarily during the adult stage, but feed and defend collectively during the larval stage in dense groups on pine branches (53–55) (Fig. 1a). When threatened by natural enemies, such as birds, ants or parasitoid wasps, the larvae perform a synchronous defensive display where they raise their head and tail (U-posture). U-posture is also required for larvae to be able to proceed to a secondary defense, where larvae release a defensive secretion and attempt to apply it directly onto the predator if it approaches close enough (56). If larvae do not lose the fluid during such encounters, they can reabsorb and reuse it during subsequent attacks (55). Previous studies have shown that sometimes this collective display alone is enough to prevent attacks by avian predators (57). Furthermore, upon the deployment of defensive fluid, volatile terpenes evaporate from the droplet, creating a repellent effect on arthropod predators such as ants and spiders (56). Additionally, the resin compounds serve as toxic irritants for arthropod predators and may cause physiological damage due to their “stickiness” (53, 56, 57)). In *N. sertifer*, survival against bird predation has shown to be higher when larvae are in a group in comparison to solitary larvae (44). Furthermore, being able to deploy the defensive fluid has been shown to improve survival in gregarious pine sawfly larvae against invertebrate predators when compared to depleted larvae (51, 58).

**Fig. 1.**
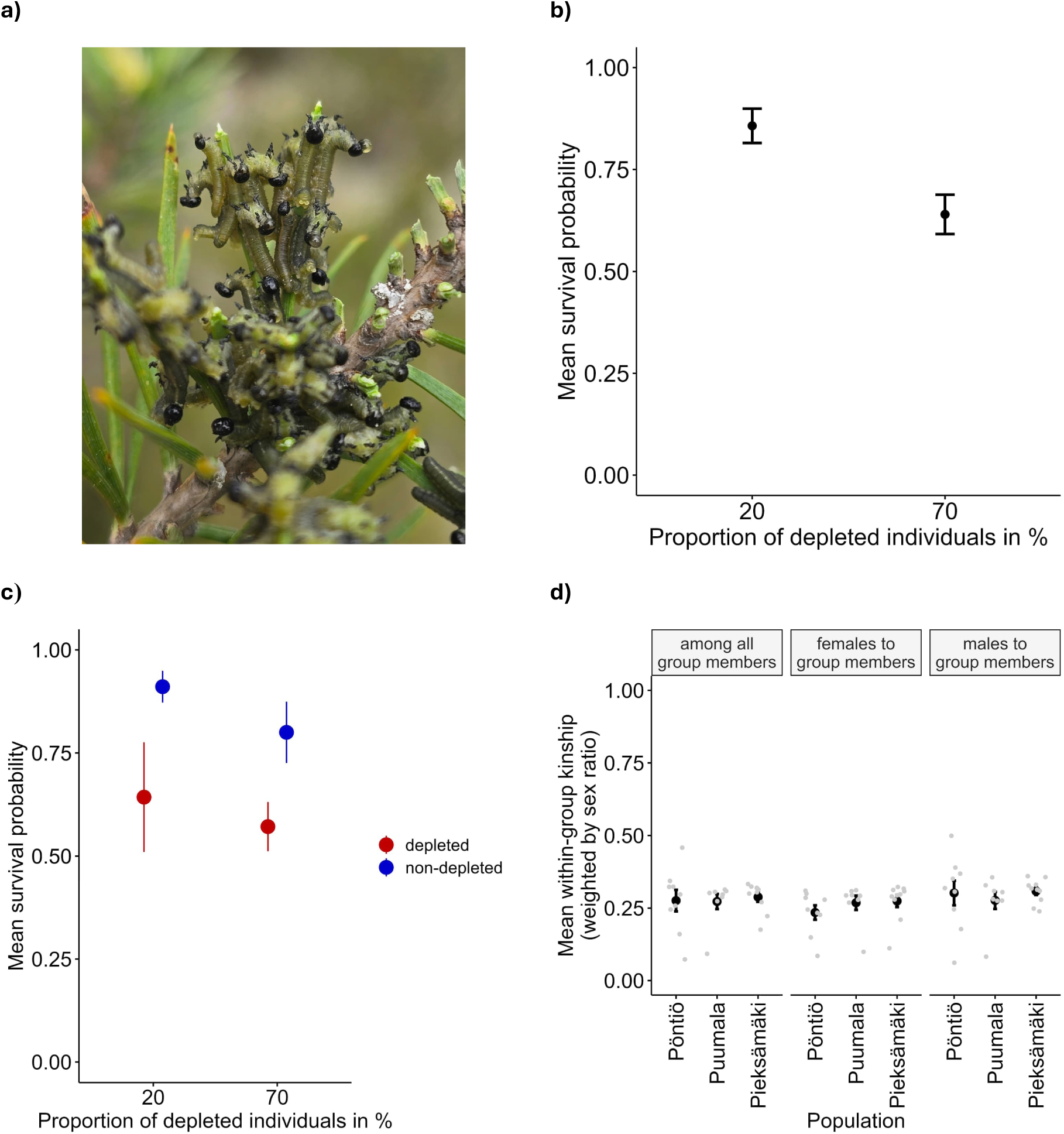
**a)** Group of mid-instar *N. sertifer* defending in the wild (photo by Raphael Ritter). **b)** Mean survival of individuals was higher in more cooperative groups (individuals in 20% depleted group: N = 70 (7 groups, 10 individuals per group), individuals in 70% depleted group: N = 100 (10 groups, 10 individuals per group). **c)** Depleted individuals (red) were less likely to survive than non-depleted individuals (blue) (depleted individuals in 20% depleted group: N = 14, non-depleted individuals in 20% depleted group: N = 56, depleted individuals in 70% depleted group: N = 70, non-depleted individuals in 70% depleted group: N = 30). **d)** Mean kinship among all group members, mean kinship of females to group members and mean kinship of males to group members of each population (in black) and larval group (in grey). Kaavi was excluded due to low sample size of males (N = 1 for mean kinship among all group members, N = 1 for mean kinship of males to other group members, and N = 2 for mean kinship of females to other group members). Error bars indicate ±1 SEM.

Pine sawfly larvae show individual and sex-biased variation in the likelihood of deploying the fluid as well as its volume and contributing to the collective chemical defense is costly (36). For example, deploying and losing the fluid repetitively decreases the growth, immune defense and life-span of individuals during the larval stage, in addition to their capacity to defend in future encounters with the predator (35, 36, 58). Interestingly, some larvae do not produce the fluid when provoked, even though their defensive reservoirs are filled, and they benefit from this via enhanced growth (36). Such non-defending individuals can be considered cheats in this system (8, 29, 36). Due to these features, it is possible to (i) quantify how much individuals are contributing to the collective defense, i.e., whether an individual deploys the fluid and the volume of fluid it produces and if so(ii) manipulate the cooperativeness within a group by manually depleting defensive fluid reserves and thereby decreasing the level of defense that can be exhibited (36). However, even when not actively secreting the defensive fluid, pine sawfly larvae remain unprofitable prey for avian and invertebrate (wood ants) predators (59–61).

We addressed five aims. First, we studied experimentally whether contribution to the collective defense provides benefits through defense against predators under natural conditions. This was done by comparing the mean survival of individuals against wood ant predation between groups containing a high proportion of defending individuals (20% of individuals depleted, 80% non-depleted) and groups containing a low proportion of defending individuals (70% of individuals depleted, 30% non-depleted). We hypothesized that individuals are more likely to survive in groups with higher proportion of defending individuals than in groups with lower proportion of defending individuals.

Second, we identified potential direct benefits of contributing to the costly collective defense. This was done by color-marking depleted and non-depleted individuals within experimental groups, and thus identifying whether depleted individuals (i.e., those that do not deploy the fluid or only in smaller amounts) are more likely to be killed by predators compared to individuals that are not depleted. We hypothesized that contribution to the collective chemical defense will be promoted if predators are less likely to attack an individual secreting the defensive fluid and/or higher volumes of it (direct benefit) (52).

Third, we investigated the importance of kin selection in maintaining defense behaviors in gregarious chemically defended prey. This was done by collecting phenotypic and genetic data from four natural populations of *N. sertifer* in Finland to quantify the kin structure and social characteristics of larval colonies and their defensive behavior. If individuals are more likely to occur in the same group with their siblings, we can expect that contribution to the costly collective defense provides indirect fitness benefits by increasing the survival of kin (16, 17). Similarly, in kin groups the relative cost of cheating increases if it decreases the survival of the cheat’s relatives (11).

Fourth, we used data gathered above to quantify the extent of variation in collective defense in natural populations. Fifth, we explored how such variation correlates with the social structure, i.e., group size, sex ratio, and kinship of groups. We hypothesized that if larval colonies vary in their relatedness structure, and if individuals were able to estimate relatedness within the group, they would be less inclined to defend (i.e., lower proportion of individuals participating in defensive U-display, deployment of fluid or its volumes) in groups with low levels of kinship. Furthermore, since females are more likely to contribute to collective defense in pine sawflies (36), we expected the mean contribution to defense to be lower when there was a higher proportion of males within a group. We also expected the individual contributions to decline with increasing group size if benefits gained by contributing to the collective defense decrease in larger groups simply due to dilution effects and individuals optimize their behavior accordingly (30, 43, 62). This local variation in the social conditions (e.g., group size (30)) could provide one mechanism maintaining differences in collective defense among populations.

## Results

### Individuals are more likely to survive predation in cooperative groups and contribution to collective defense provides direct benefits in terms of survival

The mean survival of individuals under predation by wood ants (*Formica s.str.*) was significantly higher in cooperative groups (20% depleted, i.e. non-defending individuals) compared to non-cooperative groups (70% depleted) (model 1: LRχ^2^_1_ = 5.508, estimate = -1.314, error = 0.560, p = 0.019, Fig. 1b). When the individual’s defensive status (depleted or non-depleted) was included in the model we found that depleted individuals had significantly lower survival than non-depleted individuals (model 2: LRχ^2^_1_ = 6.04, estimate = 1.932, error = 0.786, p = 0.014, Fig. 1c). However, the proportion of defending individuals within a group (model 2: LRχ^2^_1_ = 0.182, estimate = -0.331, error = 0.774, p = 0.669, Fig. 1c) or its interaction with the defensive status of an individual (model 2: LRχ^2^_1_ = 0.463, estimate = -0.652, error = 0.958, p = 0.496, Fig. 1c) did not affect survival of depleted individuals significantly (Fig. 1c). In general, the relative fitness (*v*) (11, 63) of depleted individuals was higher than or equal to that of non-depleted individuals in 4 out of 7 cases within cooperative groups, while in non-cooperative groups, depleted individuals had relatively higher survival than non-depleted ones in only 1 out of 9 cases. However, this difference is not significant and thus aligns with the results from the initial analyses (Fisher’s exact test, p = 0.105). Altogether, this suggests that depleted individuals benefit from being in cooperative groups (model 1), but that the ability to defend also benefits individuals directly (model 2).

### Contribution to collective defense provides indirect benefits

Kinship was calculated based on consanguinity values. Therefore, values for full siblings are expected to be 0.375 between sisters, 0.5 between brothers, 0.25 between sisters and brothers (64). Across all four populations, individuals show a mean kinship of 0.30 (±0.01 SEM) among females, 0.45 (±0.01 SEM) among males and 0.19 (±0.02 SEM) among females and males (Fig. 1d). When kinship estimates per group are weighted by sex ratio, mean kinship among all group members was 0.28 (±0.01 SEM), mean kinship of females to group members was 0.26 (±0.01 SEM), and mean kinship of males to all group members was 0.29 (±0.01 SEM) (Fig. S2).

### Social environment and collective chemical defense behavior vary locally

To study if the defensive behavior of *N. sertifer* larvae varied among populations, we first measured the number of individuals participating in the defensive display (i.e., U-posture) and deployment of defensive fluid per group. This was measured by attacking one randomly chosen individual in the middle of the group by poking it on the dorsal side with the tip of a pair of tweezers and calculating the number of individuals per group who displayed the U-posture or deployed the fluid.

Proportion of individuals displaying the U-posture varied significantly among populations (Table S1). This was mainly driven by the significant difference between the populations Pöntiö and Puumala as well as Pöntiö and Pieksämäki. All the other populations did not differ from each other in pairwise comparisons (Table S1, Fig. S3d). Similarly, the proportion of individuals deploying the fluid (the more costly stage of the defensive repertoire) varied significantly across populations (Table S1). Pairwise comparisons indicate that this variation was again driven by the Pöntiö population where the proportion of individuals deploying the fluid per group were lower compared to all other populations (Table S1, Fig. S3c).

Next, we analyzed how defensive behavior (whether larvae deployed the fluid or not) and the volume of fluid secreted varied among individuals and populations when larvae were attacked individually (i.e., a larva was picked up from the group and poked with tweezers). Contribution to defense was sex-biased: males were less likely to deploy the fluid than females (model 14: estimate = -1.77, error = 0.54, low 95% CI = -2.88, high 95% CI = -0.77, R-hat = 1) and produced lower volumes of it (model 15: estimate = -0.73, error = 0.21, low 95% CI = -1.13, high 95% CI = -0.31, R-hat = 1). Due to these sex biases, we analyzed the responses of females and males separately in all subsequent analyses for individually measured defense traits.

The individual’s likelihood of deploying the fluid did not vary significantly among populations, either among female or male larvae (Table S1). However, the volume of fluid individuals contributed varied significantly among populations for females (Table S1), but not for males (Table S1). In general, female larvae deployed the highest volumes in Kaavi and lowest in Pieksämäki populations (Fig. S7). Size of the individual did affect the volume deployed in males (smaller male larvae deployed higher volumes), but not in females (Table S1). In general, individuals not deploying the fluid (i.e., cheaters) were less frequent than individuals deploying the fluid (Fig. S4): across all populations, 92% of females deployed the fluid and 72% of males deployed the fluid when individually attacked.

Populations also differed significantly in terms of sex ratios and sizes of larval groups. The Pöntiö population had both the most female-biased sex ratios and largest group sizes in comparison to other populations (see Table S1 for pairwise comparisons, Fig. S3a and S3b). However, the kinship of larval colonies did not differ significantly among populations for any of the kinship types, i.e., mean kinship among all group members, mean kinship of females to group members, mean kinship of males to group members) (Table S1, Fig. 1d). We excluded Kaavi from the analyses because of low sample size due to the lack of males (N = 1 for mean kinship among all group members, N = 1 for mean kinship of males to other group members, and N = 2 for mean kinship of females to other group members). Similarly, the kinship of larval colonies did not differ significantly among populations for female-female, male-male or female-male kinship estimates which were not weighted by the sex ratio of the group. (Table S4).

### Individuals regulated their contribution to the collective chemical defense based on their social environment

The mean kinship of females to group members was significantly positively correlated with the display of the U-posture. The other kinship types did not correlate significantly with the display of the U-posture (Table 1, Fig. 2c). Neither sex ratio nor group size had a significant effect for the proportion of individuals which displayed U-posture synchronously when provoked (i.e., one individual per group attacked), (Table 1, Fig. S5a and S5b).

**Table 1.** Effect of sex ratio, group size and kinship on the proportion of individuals per group performing the U-posture or deploying the fluid. Significant values are in bold (p < 0.05).

| Model # | Fixed factor | Chi<sup>2</sup> | Beta | Error | p-value |
| --- | --- | --- | --- | --- | --- |
| Proportion of individuals displaying the U-posture within a group |  |  |  |  |  |
| 16 | Sex ratio | LR $\chi^2_1 = 2.002$ | -1.379 | 0.974 | 0.157 |
| | Group size | LR $\chi^2_1 = 0.378$ | -0.005 | 0.008 | 0.539 |
| 17 | Mean kinship among all group members | LR $\chi^2_1 = 0.480$ | 2.065 | 2.981 | 0.489 |
| 18 | Mean kinship of females to group members | LR $\chi^2_1 = 4.103$ | 9.034 | 4.462 | **0.043** |
| 19 | Mean kinship of males to group members | LR $\chi^2_1 = 0.010$ | 0.272 | 2.660 | 0.919 |
| Proportion of individuals deploying the defensive fluid within a group |  |  |  |  |  |
| 20 | Sex ratio | LR $\chi^2_1 = 6.564$ | -1.428 | 0.557 | **0.010** |
| | Group size | LR $\chi^2_1 = 13.766$ | -0.015 | 0.004 | **<0.001** |
| 21 | Mean kinship among all group members | LR $\chi^2_1 = 0.744$ | 1.296 | 1.503 | 0.389 |
| 22 | Mean kinship of females to group members | LR $\chi^2_1 = 1.501$ | 2.022 | 1.650 | 0.221 |
| 23 | Mean kinship of males to group members | LR $\chi^2_1 = 0.912$ | 1.390 | 1.456 | 0.340 |

**Fig. 2.**
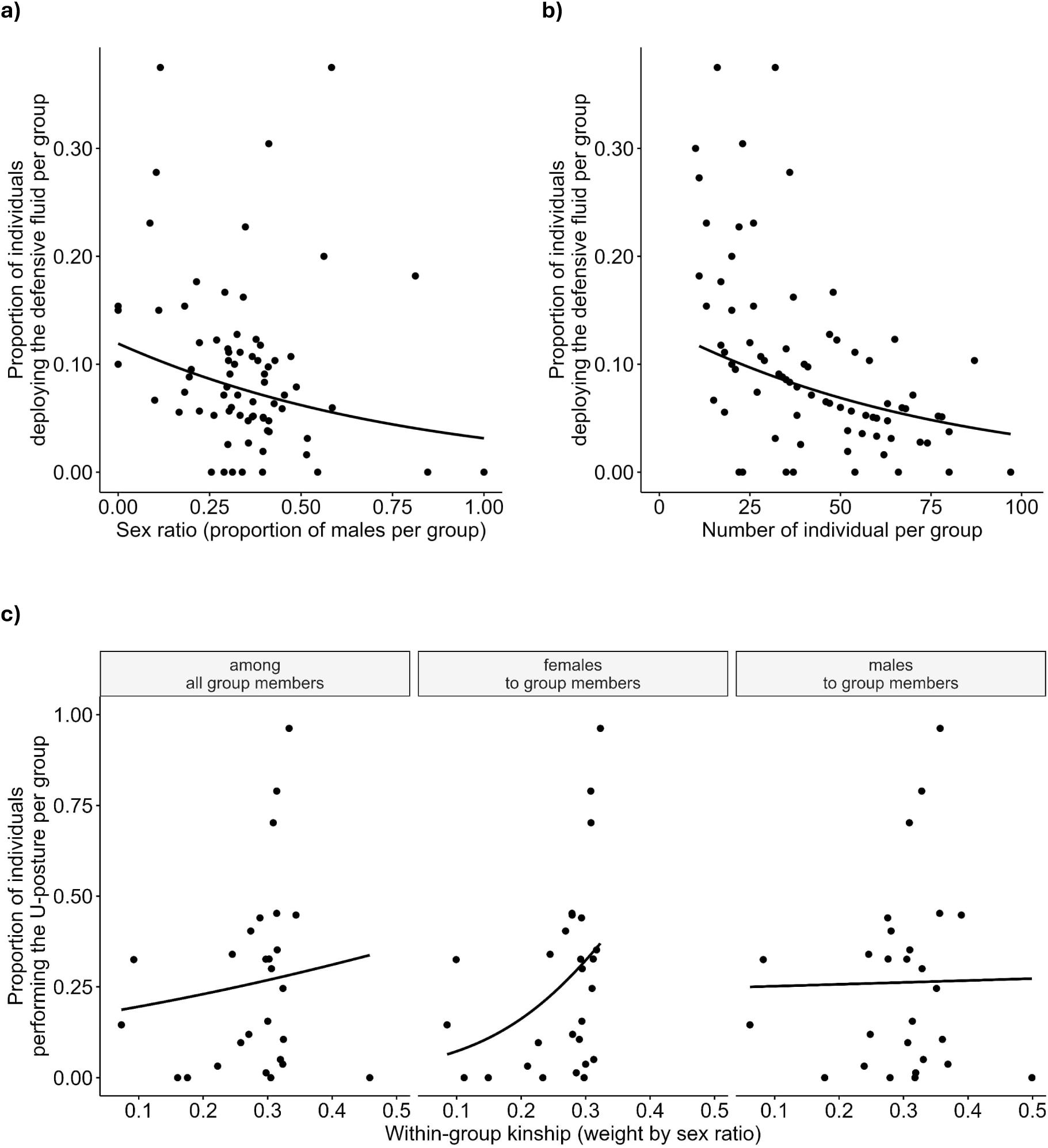
Phenotypic association between the contribution to the collective defense (secretion of defensive fluid) and the **a)** sex ratio (proportion of males) (N = 74) and **b)** size of the larval group (N = 83). **c)** Phenotypic association between participation in the defensive display (U-posture) and the kinship level of larval groups (N = 27 for each kinship type).

When the proportion of individuals per group deploying the fluid was analyzed, cooperativeness of the group decreased significantly with an increasing proportion of males. Similarly, proportion of individuals that deployed the fluid decreased with the increase in group size (Table 1, Fig. 2a and 2b). However, none of the kinship types were correlated with the proportion of individuals deploying the fluid within a group (Table 1, Fig. S5c).

Similar analyses were run for the individually measured defensive traits. Female larvae deployed significantly higher volumes of fluid with higher level of kinship within a group (mean kinship among all group members, mean kinship of females to group members, mean kinship of males to group members) (Table 2, Fig. S6). Group size and sex ratio did not influence the volume of fluid secreted (Table 2). For males, neither sex ratio, group size nor any of the kinship types affected their probability of deploying defensive fluid or the amount of fluid secreted (Table 2). This was also the case for females’ probability of deploying the defensive fluid. Larval length did not affect how much fluid was deployed in both females and males in these models.

**Table 2.**
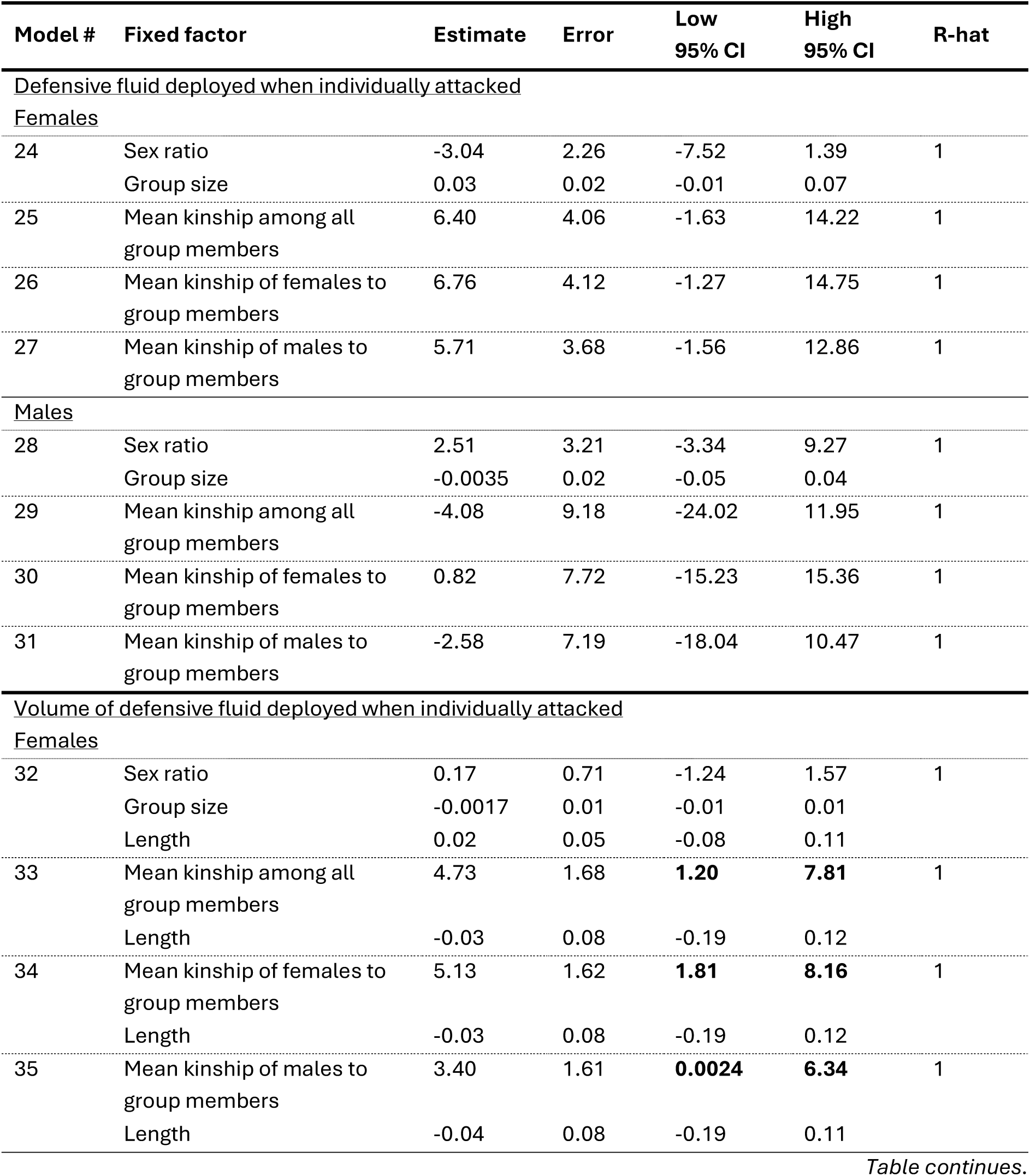
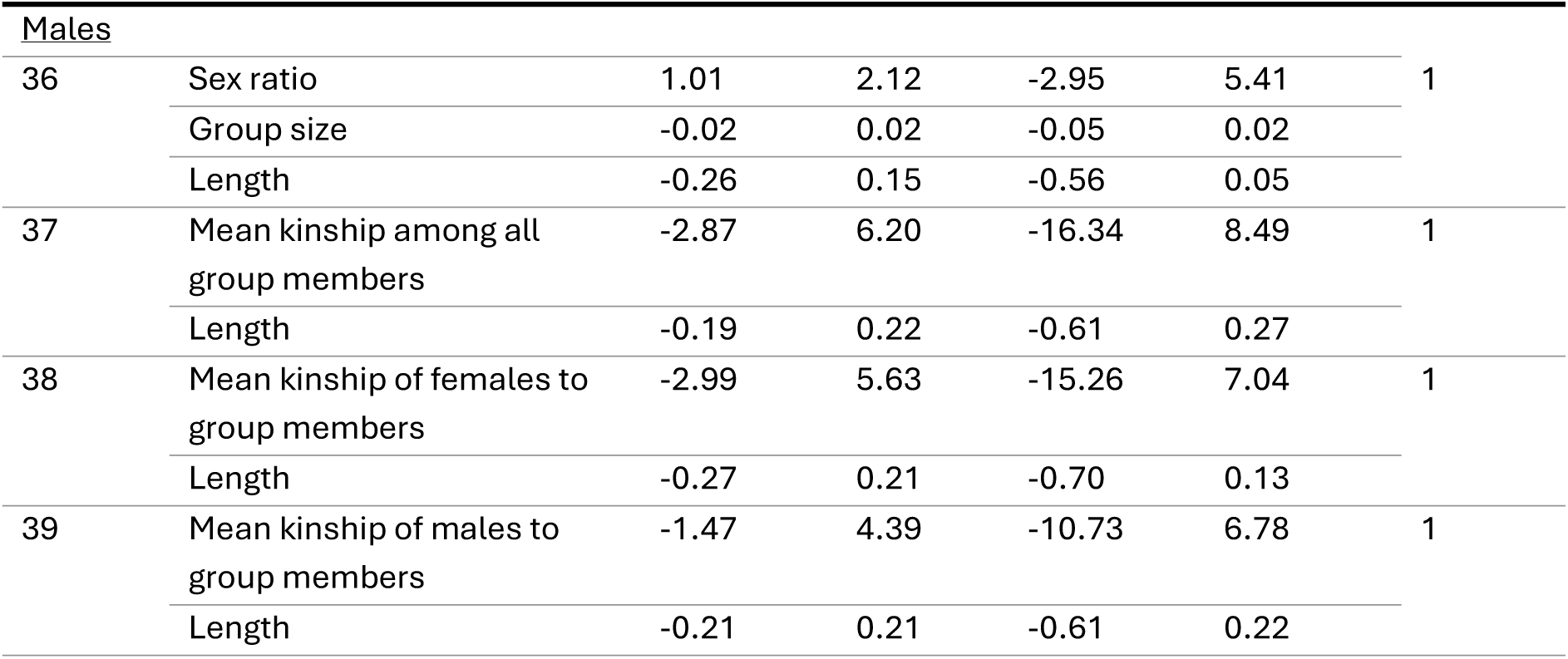
Effect of sex ratio, group size, and kinship on individuals’ defensive traits. Variables whose 95% credible intervals do not include zero are in bold.

| Model # | Fixed factor | Estimate | Error | Low<br>95% CI | High<br>95% CI | R-hat |
| --- | --- | --- | --- | --- | --- | --- |
| <u>Defensive fluid deployed when individually attacked</u> |  |  |  |  |  |  |
| <u>Females</u> |  |  |  |  |  |  |
| 24 | Sex ratio | -3.04 | 2.26 | -7.52 | 1.39 | 1 |
|  | Group size | 0.03 | 0.02 | -0.01 | 0.07 |  |
| 25 | Mean kinship among all<br>group members | 6.40 | 4.06 | -1.63 | 14.22 | 1 |
| 26 | Mean kinship of females to<br>group members | 6.76 | 4.12 | -1.27 | 14.75 | 1 |
| 27 | Mean kinship of males to<br>group members | 5.71 | 3.68 | -1.56 | 12.86 | 1 |
| <u>Males</u> |  |  |  |  |  |  |
| 28 | Sex ratio | 2.51 | 3.21 | -3.34 | 9.27 | 1 |
|  | Group size | -0.0035 | 0.02 | -0.05 | 0.04 |  |
| 29 | Mean kinship among all<br>group members | -4.08 | 9.18 | -24.02 | 11.95 | 1 |
| 30 | Mean kinship of females to<br>group members | 0.82 | 7.72 | -15.23 | 15.36 | 1 |
| 31 | Mean kinship of males to<br>group members | -2.58 | 7.19 | -18.04 | 10.47 | 1 |
| <u>Volume of defensive fluid deployed when individually attacked</u> |  |  |  |  |  |  |
| <u>Females</u> |  |  |  |  |  |  |
| 32 | Sex ratio | 0.17 | 0.71 | -1.24 | 1.57 | 1 |
|  | Group size | -0.0017 | 0.01 | -0.01 | 0.01 |  |
|  | Length | 0.02 | 0.05 | -0.08 | 0.11 |  |
| 33 | Mean kinship among all<br>group members | 4.73 | 1.68 | **1.20** | **7.81** | 1 |
|  | Length | -0.03 | 0.08 | -0.19 | 0.12 |  |
| 34 | Mean kinship of females to<br>group members | 5.13 | 1.62 | **1.81** | **8.16** | 1 |
|  | Length | -0.03 | 0.08 | -0.19 | 0.12 |  |
| 35 | Mean kinship of males to<br>group members | 3.40 | 1.61 | **0.0024** | **6.34** | 1 |
|  | Length | -0.04 | 0.08 | -0.19 | 0.11 |  |
| <u>Males</u> |  |  |  |  |  |  |
| 36 | Sex ratio | 1.01 | 2.12 | -2.95 | 5.41 | 1 |
|  | Group size | -0.02 | 0.02 | -0.05 | 0.02 |  |
|  | Length | -0.26 | 0.15 | -0.56 | 0.05 |  |
| 37 | Mean kinship among all group members | -2.87 | 6.20 | -16.34 | 8.49 | 1 |
|  | Length | -0.19 | 0.22 | -0.61 | 0.27 |  |
| 38 | Mean kinship of females to group members | -2.99 | 5.63 | -15.26 | 7.04 | 1 |
|  | Length | -0.27 | 0.21 | -0.70 | 0.13 |  |
| 39 | Mean kinship of males to group members | -1.47 | 4.39 | -10.73 | 6.78 | 1 |
|  | Length | -0.21 | 0.21 | -0.61 | 0.22 |  |

When we analyzed how social characteristics of larval groups were correlated, we found that female-female and female-male kinship decreased in larger groups. (Table S3, Fig. S8c). That is, in larger groups, individuals were less related. The sex ratio of the group was not associated with group size or any of the kinship estimates (Table S3, Fig. S8a and S8b).

Finally, populations were not genetically structured. First, we found that F_ST_ values were close to zero for all pairwise population comparisons and all female combinations (Fig. S9a). Second, F_IS_ values were slightly negative for most combinations of females (Fig. S9b). These results suggest panmixia, i.e., low differentiation across large distances and indicate lack of inbreeding within populations.

## Discussion

Determining factors that drive cooperation and cheating in public goods is fundamental to predict variation in cooperative interactions within and between species (24). Based on Hamilton’s inclusive fitness theory, we can expect selection to favor cooperation when the direct and indirect fitness benefits to the actor outweigh the fitness cost. Here, we have demonstrated experimentally that collective chemical defense is beneficial through improving survival of individuals, and that this results in indirect benefits as most individuals within larval colonies are full siblings. We have also shown that such cooperation confers direct benefits, as individuals that had maintained their capacity to defend had a higher probability of survival than those that lost it. Our phenotypic data from wild groups reveal significant variation in collective defense behavior across groups and populations. This variation among larval groups is partly explained by social context, with individuals becoming less inclined to defend in larger, more male-biased groups, as well as when the level of kinship decreases. Contribution to collective chemical defense was also sex-biased, with females more likely to deploy the defensive fluid and to do so in greater volumes than males, especially when they are closely related to other group members. Together, these results suggest that while contributing to defense improves individual survival, individuals adjust their contributions to the collective good based on their social environment and sex. Variation in these factors leads to within- and among-population variation in cooperative behavior.

Public goods systems are often vulnerable to cheating, but cheats might not outcompete cooperative phenotypes completely if cooperation has high direct benefits. Our data from natural populations indicates that cheating in *N. sertifer* populations is rare among females but not unusual among males (8% of females and 28% of male larvae do not defend when individually attacked and measured; Fig. S4). However, we did not find strong support for the frequency dependent benefits of cheating (11) (experimentally represented by depleted individuals). For example, while the relative fitness (11, 63) of depleted individuals was more often lower when they were common (8/9 cases in 70% depleted) compared to when they were rare (3/7 cases in 20% depleted), this difference was not significant (see also Fig. 1b and 1c). This suggests that at least under high predation intensity, contribution to collective defense is promoted via high direct benefits of defense. It remains possible that frequency-dependent costs and benefits could occur at more extreme frequencies not included into our experimental design. Additionally, our experiment included only one type of predator (wood ants). In nature, predator species vary in their sensitivity to defense mechanisms of prey (reviewed in (65)) and in their responses to visual cues and signals of prey individual’s unprofitability (66, 67). Therefore, the benefits of contributing to group defense can vary depending on the intensity of predation as well as predator community structure (65–68). While we used a natural predator in our experiments, how cooperation is selected under diverse predation pressures is clearly an interesting avenue for future research.

The benefits of cooperation in public goods can also depend on group size (30). For example, studies on public goods systems in microbes have shown that the fitness of cheats increases at high population densities (62). Accordingly, our phenotypic data from natural populations suggest that the proportion of individuals contributing to collective defense decreases as group size increases. At larger aggregations, it might be more beneficial for a gregarious prey to rely on the dilution effect and allocate energy and resources to growth instead (32, 33, 43, 69). Group sizes also varied quite extensively within and among populations (Fig. S3b). Part of this variation can be a result of merging of offspring from several females (51, 70). This is supported by our observation that the within-group kinship decreases in larger group sizes. In addition, the quality of the host plant can affect both oviposition decision of females (71, 72) as well as the hatching success of eggs (71) and mortality rates of larvae (73), further influencing larval group sizes.

Our results show that similarly to *Diprion pini* (36), also *N. sertifer* females contribute to collective defense significantly more often and in greater quantities than males. In addition, both the sex ratios of populations and the larval colonies were female-biased (Fig. S3). The recent theoretical model focusing on the pine sawfly system (74) showed that when sex ratios are female-biased during the larval stage, female-biased helping in collective defence can evolve due to the rarer-sex effect, even when defensive costs would be higher for females than for males (74). It is also possible that pine sawfly females are pre-adapted to help, given that they grow larger than males in the larval stage and are thus able to contribute to defending more effectively. This is similar to termites, where size could be a pre-adaptation promoting the sex biased division of labor among females and males (75). More generally, maternal behaviors as pre-adaptations (rather than haplodiploidy *per se*) explain the prominence of female helpers in social Hymenoptera (76, 77). The evolution of sex-biased helping can also be linked to haplodiploidy and the easy sex ratio adjustments it allows, given that sex of the eggs is determined by fertilization (78). As mothers may be under selection to produce the more helpful sex (i.e., local resource enhancement), and the reproductive value is lower for the individuals of the more common sex (“the rarer sex effect”) (79, 80) female-bias may drive a positive feedback loop between sex ratio bias and sex-biased helping (78). Given the female-bias in *N. sertifer* larval groups and populations (see also (81–83) for female-biases in other sawfly species), and that sex-biased allocation to the collective defense results in a higher level of chemical defense at the group level (Fig. 2a), this coevolutionary feedback loop seems a promising topic for future research.

Kin selection plays an important role in promoting cooperation across many different species (84–86). Similarly, our results show that pine sawfly larvae occur in kin groups, providing indirect benefits to the defending individuals. Within-group kinship values suggest that single matriline groups where the mother is singly mated are the rule, but occasional all male groups from unmated females, multiple mating or multiple matriline groups may occur (82). Facultative adjustment of cooperativeness based on the variation in kinship was observed in females, further highlighting the importance of indirect benefits. Thus, while the clear direct benefits of defense may imply that the benefit to kin is not the main driver of defending behavior, the existence of an indirect fitness benefit suggests that the trait is cooperative in nature, as it evolves at least partially due to the benefits it provides to others (87). Nevertheless, to better understand the cause-and-effect relationships between kinship, group size, and sex ratio on the contribution to collective chemical defense, controlled experiments are required in which their effects are manipulated and measured independently of each other.

Finally, our results also provide new insights into the evolution of costly chemical defenses. Although R. A. Fisher suggested already in 1930 (79) that kin aggregations could promote the evolution of prey distastefulness, the importance of kin selection in favoring the evolution of costly chemical defenses has received much less attention (40) compared to the theoretical and empirical work on the direct benefits it provides (43, 45, 88). Our results here suggest that kin aggregations can play an important role and should be considered when examining the early evolution of costly antipredator defenses. However, in the model first proposed by Fisher (79) and later developed by Leimar et al. (88) and Best et al. (40) the chemical defenses were stored within the prey, and there were no cues or signals indicating their distastefulness to predators. This means that predators need to sample, taste and kill the prey to identify its defensive status before learning to avoid it. Our findings suggest that for gregarious prey that excrete defensive secretions externally, the direct benefits could be sufficient to promote the evolution of these traits, as defended prey do not necessarily all die before local predators learn to avoid them.

More generally, we demonstrate that ecological drivers of cooperation can play an important role in its maintenance, and that the facultative expression of cooperativeness due to social conditions can significantly contribute to the maintenance of local variation in its expression.

## Material and methods

### Predation experiment with ants

The predation experiment with ants was conducted in the summer 2023. The larval families used in this experiment originated from insect cultures established from the wild individuals originating from the larval colonies collected in different parts of Finland. Separate insect cultures per population were established with the following number of colonies collected from the wild: Alavus (50 colonies in 2020), Kaavi (10 colonies 2021 and 40 colonies 2022), Pieksämäki (40 colonies 2021 and 20 colonies 2022), Pöntiö (20 colonies 2021), Seinäjoki (10 colonies 2021). Experimental individuals originated from F1-F3 generations depending on the starting year of the insect culture. They were reared in a laboratory at the University of Helsinki, where they were kept at room temperature and fed fresh pine needles *ad libitum*. At approximately 10 days of age, larval groups were assigned to the depletion treatment or left to retain their defensive fluid. Depleted and non-depleted larvae were identified with fluorescent elastomer tags (89, 90). These were injected subcutaneously and function well in pine sawfly larvae as they remain through instars. We used either fluorescent orange or blue dye (Visible Implant Elastomer Tags by Northwest Marine Technology, Inc.) and varied the colors for depleted and non-depleted treatment across groups to avoid the color markings to confound the effects of the experimental treatments. On the same day, we simulated an attack by gently pressing them with a cotton swab on the dorsal side. If they regurgitated defensive fluid, it was swiftly removed with the same cotton swab. Our supplementary data (Fig. S10) suggests that this procedure should be repeated multiple times to ensure that most of the depleted larvae can no longer deploy the defensive fluid. We therefore repeated this once more, two days later. Larvae were then placed in groups of 10 on a fresh pine twig, mixing depleted and non-depleted sibling individuals to create groups with 20% or 70% depleted individuals per group. The groups were then left undisturbed until the next morning. We did not mix individuals of different families, i.e., larvae were presented to ants in full-sib groups.

In the field, we placed an approximately 40 cm long pine branch into the ground on a foraging trail in the vicinity of *Formica s. str.* mound and attached the twig bearing the group of larvae to it using green coated metal wire. Such replicates were carried out at 8 different ant nests. Altogether, we had 7 replicates for the 20% treatment and 10 replicates for the 70% treatment. We always had all the treatment combinations within the same nest, except for two nests where we only had one of the two treatment groups. We recorded the survival of the larvae over a seven-hour period.

### Collection of *N. sertifer* specimen for measurements of defense behaviors and social environmental factors

We collected larvae in summer 2021 from different parts of Finland (Pöntiö, Pieksämäki, Puumala, and Kaavi) with only one group taken from each tree per study site. Each group consisted of larvae found in close proximity to one another on the same branch of the tree, clearly forming a single group. These groups were subsequently reared in a greenhouse at the University of Jyväskylä, where the temperature matched natural outdoor conditions. Larvae were fed fresh pine branches ad libitum. Lighting conditions always included both natural sunlight and some level of electric light. To measure the defensive behavior of each group, we simply randomly selected one individual per group, poked it gently (non-injuriously) on its dorsal side with the tip of a pair tweezers, and counted the number of larvae that responded by regurgitating defensive fluid and/or raising their heads in response to the simulated attack. When larvae formed subgroups within each aggregation, we simulated an attack on one individual randomly within each subgroup. We measured a total of 84 groups (Kaavi: N = 8, Pöntiö: N = 20, Puumala: N = 18, Pieksämäki: N = 38). We determined the size of each larval group immediately after these measurements. We sexed the individual larvae based on the adults that eventually emerged and when an individual pupated but failed to eclose, we used cocoon size as an estimator for sex (females being larger than males (36, 55)). The sex of 70 individuals were unknown as they died before pupation. For the individual defense measurements, we simply randomly chose 5 individuals from each group and simulated an attack on each one. We also sucked any fluid secreted into a glass capillary and measured the length of the tube occupied by the liquid (using calipers accurate to 0.1 mm) and converted the measure to a volume (mm^3^). In total, we measured the defense behavior of 345 individuals from 69 different groups (Kaavi: 10 groups, 50 individuals, Pöntiö: 20 groups, 100 individuals, Puumala: 19 groups, 95 individuals, Pieksämäki: 20 groups, 100 individuals).

### DNA extraction s preparations for sequencing

The majority of groups had at least 10 females and males, but due to mortality and variation in original larval group sizes and sex-ratios, some contained only of 5 females or 1 male per larval group. We included only groups that contained both females and males. We followed the Qiagen kit instruction with minor modifications. After extraction, we normalized the DNA concentration to approximately 20ng/µL and double-digested it with the enzymes Mspl and MluCl. To associate each sequence with a specific 96-well plate, we ligated double-stranded adapters to each DNA strand. We then conducted a test-PCR, followed by gel electrophoresis to confirm successful DNA digestion and adapter ligation, which led to the exclusion of 14 samples. The remaining samples were pooled into a single 96-well plate and cleaned using an AMPure substitute. Next, we attached index primers with unique dual indices (UDI) to associate each sequence with a specific position on the 96-well plate. All samples were then pooled into a single Eppendorf tube and run on a 1.5% agarose gel for size selection (320-500 kb). We excised the DNA region from the agarose gel and purified it with the Monarch DNA Gel Extraction Kit (New England Biolabs). The cleaned library was quantified via qPCR using the NEBNext Library Quant Kit (New England Biolabs) and sequenced on the Illumina NovaSeq platform at the Institute for Molecular Medicine Finland (FIMM) of the University of Helsinki. Altogether, after the filtering steps described below, we sampled from Pöntiö 10 groups and 189 individuals, from Pieksämäki 10 groups and 186 individuals, from Puumala 9 groups and 166 individuals, and from Kaavi 8 groups and 69 individuals.

### Bioinformatics (kinship analysis)

We trimmed the 4bpIMI off the raw sequencing data from the left of each forward read and then demultiplexed them based on the 5bp index allowing us to identify the sample name for each read. Following this, we used the ipyrad pipeline to assemble RAD loci for each individual. For the parameter settings, we deviated from the default settings as follows: Assembly method: pairddrad, restriction overhang: AATT, GC, filter_adapters: 2. We then filtered the vcf file using *vcftools* (91) with the following arguments: --maf 0.01, --remove-indels, --max-alleles 2, --max-missing 0.5. We further retained only one random SNP per locus using the python script vcf_parser.py (https://github.com/CoBiG2/RAD_Tools/blob/master/vcf_parser.py). Pairwise kinship, i.e., consanguinity, was calculated using the R package *HapDipKinship* (64). We encoded male haploid genotypes as 0 (reference allele) or 1 (alternative allele). Any heterozygous genotype calls in haploids were treated as errors and set to missing (NA). Female diploid genotypes were encoded as 0 (homozygous reference), 1 (heterozygous), and 2 (homozygous alternative).

### Bioinformatics (population genetic data)

For the population genetics, we randomly chose one female per group from each population, resulting in a sample size of 37 individuals, and ran the analysis for 20 different combinations of females. To infer the population structure, we performed a clustering analysis with ADMIXTURE (92), testing for different number of clusters/populations (*K*) ranging from 1 to 10. We determined the most suitable value for *K* by using ADMIXTURE’s cross-validation (CV) procedure, comparing the CV error values for each *K*. We further checked for population differentiation and inbreeding by calculating F_ST_ and F_IS_ values using the R packages *adegenet* (93, 94) and *hierfstat* (95), respectively.

### Statistical analysis

We performed all statistical analyzes using R version 4.2.2 (R Core Team, 2024). We used *glmmTMB* v1.1.10 (96) to run all Frequentist models and the *DHARMa* package v0.4.7 (97) with the *simulateResiduals* function to test and confirm the model assumptions. When we could not confirm Frequentist model assumptions we constructed Bayesian models using the *brm* function of the *brms*package, v.2.22.0 (98–100). For all models, we used the default priors provided by the *brm* function, i.e., flat or weakly informative priors for the intercept, fixed and random effects. We checked for convergence and autocorrelation using the R-hat values as well as assessed visually trace and autocorrelation plots. All R-hat values were at 1, autocorrelation plots showed rapid decay, and trace plots indicated stable patterns. There were divergent transitions for the following modes: model 33 (13 transitions), 35 (3 transitions), 37 (1 transition), 40 (2 transitions), 41 (1 transition), 42 (14 transitions). We checked model fits using pareto k diagnostics with the *loo* package 2.8.0 (101–103). Influential points (k > 0.7) were detected for model S3*(1count), 15 (1 count), 26 (2 counts), 29 (2 counts), 30 (2 counts), 31 (2 counts), 36 (2 counts), 37 (1 count), 38 (1 count), 39 (2 counts), S3 (1 count), S11 (1 count), S12 (2 counts), S13 (2 counts), S15 (1 count), S16 (1 count), S17 (2 counts). Model specifications of each model are listed in Table S8. We obtained beta coefficients and their standard errors from the model summary, and chi-squared values and p-values from the Anova function of the *car* package, v3.1-3 (104).

We centered all numeric explanatory variables (i.e., predictors have a mean of zero) to improve interpretability of the results. For those models that included one of the kinship types and/or male individual traits (model 6, 8, S2, S3, S3*, 9, 9*, 10, 10*, 14, 15, S5, S6, 17, 17*, 18, 19, S8, S9, 21, 22, 23, S11, 25, 26, 27, 28, S12, S13, 29, 30, 31, S15, 33, 34, 35, 36, S16, S17, 37, 38, 39, 42, 43), we excluded the Kaavi population because of low sample size (N = 2 for female-male comparisons and N = 1 for male-male comparisons, N = 2 for individual-level traits). Sex ratio was calculated of groups that consisted of at least 10 larvae of known sex. We excluded 25 pairwise kinship estimates with large negative values ranging from -8 to -Inf. All data wrangling and visualizations were done with the *tidyverse* v2.2.0 (105) and *ggpubr*v0.6.0 package (106). We calculated the mean kinship of females and males to their group members as well as overall kinship of all group members as follows:

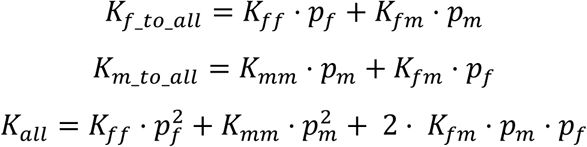

K_all_: mean kinship among all group members

K_f_to_all_: mean kinship of females to group members

K_m_to_all_: mean kinship of males to group members

K_ff_: mean kinship among females within a group

K_mm_: mean kinship among males within a group

K_mm_: mean kinship among females and males within a group

p_f_: proportion of females within a group

p_m_: proportion of males within a group

We constructed separate models for each of these kinship types (i.e., mean kinship among all group members, mean kinship of females to other members, mean kinship of males to group members). As *glmmTMB* and *brm* are only able to make use of those lines that have data for all variables included in the model, using all 5 variables in the same model would have led to loss of the majority of sex ratio and group size data.

### Predation experiment with ants

We constructed generalized linear mixed models with a binomial distribution and a logit link function. In the first model, we examined whether survival differs between groups with lower (20%) and higher (70%) proportions of depleted individuals. We used individual survival at the end of the experiment as the binary response variable and included the proportion of depleted individuals as a fixed factor. To account for variation between ant nests (e.g., in number of individuals, activity and behavior), we included Ant nest ID as a random factor. To account for potential effects of shared environment (i.e., larval group and/or location of larvae on the ant trail) on survival, we additionally included Group ID as random factor. In the second model, testing for individual benefits of contributing to the defense, we included an interaction term between depletion treatment and the proportion of depleted individuals as fixed factor. We also included Ant nest ID and Group ID as a random effect. The relative fitness *v* of depleted individuals for each group was calculated based on (63):

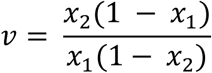

where *x_1_* is the initial proportion of cheats (i.e., depleted individuals) within a group (0.7 in 70% treatment and 0.2 in 20% treatment) and *x_2_* is their final proportion (number of depleted individuals survived/group size at the end of the experiment). Groups were then divided into two categories based on the *v*-value: groups where *v* < 1 belonged to category 0 and groups where *v* = 1 or > 1 (i.e., relative fitness of depleted individuals was equal or higher to non-depleted ones) belonged to category 1. One of the groups where only one individual survived was excluded from these analyses due to issues with “nullity” (i.e., not possible to divide values by zero). We used Fisher’s exact test to analyze if there is a significant association between the 20% and 70% treatment and relative fitness of cheats.

### Phenotypic and kinship data

#### U-posture display within a group

To test how displaying the U-posture in response to a simulated attack varies with the social environment, we used generalized linear mixed models. We modeled the response variable as binomial data. We constructed separate models for each kinship type (mean kinship among all group members, mean kinship of females to group members, mean kinship of males to group members) as well as group size and sex ratio to analyze how each of them impacts whether individual larvae display the U-posture. In all four models, we included Population ID as a random effect to account for variation among populations in the display of the U-posture. As our data was overdispersed, we used a beta-binomial distribution with a logit link function.

#### Deployment of defensive fluid within a group

To test how cooperative behavior (deploying the fluid) varies with the social environment, we used the same approach as when analyzing the U-posture with the difference that we used a binomial distribution as data was not overdispersed.

#### Differences between sexes

To analyze whether females and males differ in their defense behavior, we used Bayesian modelling. One model for the measure of fluid secreted or not, and one for the measure of the volume of fluid secreted. In both cases, we used sex as a fixed factor, and Group ID was nested under Population ID as a random factor to account for potential genetic and phenotypic variation among groups and populations in cooperativeness. For the binary data (fluid deployed or not), we used a binomial distribution with logit link function. For the volume data, we used a gamma distribution because of the strongly right skewed distribution of the response variable with a log link function. Because gamma distributions do not allow for zero values, two decimals more than the precision of the data (i.e., 0.001) was added to the values of the defensive fluid in all the analyses for the volume of defensive fluid (see also below). We included also those individuals that did not regurgitate any fluid in this analysis to take into account the whole extent of the observed variation.

#### Individual defense and social environment

We used Bayesian modelling to analyze whether the social environment of the larvae affects the defense behavior (deployment of fluid and its volume) when individually attacked. Due to the sex-biased defense behavior (36), we analyzed females and males separately. We constructed separate models for each kinship type (i.e., mean kinship among all group members, mean kinship of females to group members, mean kinship of males to group members) to test whether group size, sex ratio and kinship impacts the individual defense. In all 16 models, we included Population ID and Group ID as a random effect to account for potential differences among populations and groups. In the 8 models analyzing whether individuals secreted the fluid or not, we used a binomial distribution with a logit link function. In the other 8 models analyzing the amount of fluid that individuals regurgitated, we used a gamma distribution because of the strong right skewed distribution of the response variable with a log link function. To account for allometric effects, we additionally included larval length as a fixed factor. We included also those individuals that did not regurgitate any fluid in this analysis.

#### Local variation of social environment and collective chemical defense behavior

To analyze whether populations differ in cooperativeness and social environmental factors, we constructed 11 models, one for each trait: proportion of individuals deploying the fluid, proportion of individuals displaying the U-posture, individual defense (fluid deployed or not) of females, individual defense (fluid deployed or not) of males, individual defense (defensive fluid volume) of females, individual defense (defensive fluid volume) of males, group size, sex ratio, mean kinship among all group members, mean kinship of females to group members, mean kinship of males to group members. For traits that were measured on the individual level (i.e., fluid deployment of females and males, volume of fluid produced by females and males), Group ID was added to the models as a random effect. Larval length was additionally included when modelling the amount of defensive fluid deployed. For the binary data (deployed the fluid or not), we used a binomial distribution with a logit link function. Sex ratio, group size, and the kinship data were approximately normally distributed and we therefore modelled it with a gaussian distribution and identity link function. Secreted defense volume was strongly right skewed and contained zeros. We therefore used a tweedie distribution with a log link function. As we failed to confirm the model assumptions of model 9 (mean kinship among all group members) and model 10 (mean kinship of females to group members), we re-ran them results with a Bayesian approach which confirmed the results we received from the Frequentist approach.

#### Correlations between traits

To test correlations between the social characteristics of groups, we applied a Bayesian modelling approach. To test how group size changes with the sex ratio of the group, we used the proportion of males as the fixed factor and Population ID as a random factor, accounting for potential within-population similarities. To test how kinship is affected by group size and sex ratio, we constructed a model for each kinship type with Population ID as the random factor. We used a gaussian distribution with identity link function. Here, we used kinship estimates unweighted by sex ratio as we included sex ratio as a fixed factor. We therefore ran three separate models with female-female kinship, male-male kinship and female-male kinship.

## ACKNOWLEDGEMENTS

This study was funded by the Research Council of Finland via the project no 330578 (CL) and 333482 (SW). We are grateful to Nina Immonen, Liina-Lyydia Jämsä, Venla Korhonen, and Jenni Sohlman for helping with the maintenance of lab population and collecting field data as well as local Forest Centers in Central Finland for providing information about the potential outbreak areas.

Authors declare no conflict of interests.

## AUTHOR CONTRIBUTION STATEMENT

CL conceived the ideas and CL, RR, RT, HH and SW designed methodology; RR and RT collected the data; RR, RT and SW conducted genetic and phenotypic analyses of the data; RR led the writing of the manuscript, with input from CL, HH, and SW. All authors contributed critically to the drafts and writing and gave final approval for publication.

## DATA AVAILABILITY STATEMENT

All data sets and scripts used in these analyses are archived in the open access database Dryad (https://doi.org/10.5061/dryad.ngf1vhj85).

