## Supplementary material for "Direct and indirect benefits of cooperation in collective defense against predation": SI Text

a) Department of Forest Sciences, University of Helsinki, University of Helsinki 00014, Finland
b) Ecology and Genetics research unit, University of Oulu, Oulu 90014, Finland
c) Natural Resources Institute Finland (Luke), Joensuu 80100, Finland
d) Department of Environmental and Biological Sciences, University of Eastern Finland, Joensuu 80101, Finland

**This file includes:**

SI Text, Figures S1 to S10, Tables S1 to S8, SI References

**Other supporting materials for this manuscript include the following datasets:**

S1: Data set: Predation experiment

S2: Data set: Group level measurements

S3: Data set: Individual level measurements (females and males)

S4: Data set: Individual level measurements (females only)

S5: Data set: Individual level measurements (males only)

S6: Data set: Pairwise kinship estimates

S7: CV errors of population genetic analysis

S8: $F_{IS}$ values of population genetic analysis

S9: $F_{ST}$ values of population genetic analysis

S10: Data set: Effect of depletion treatment on defensive fluid deployment

S11: R code of all models and plots presented

S12: Data set and codes: Kinship analysis

S13: Data set and codes: Population structure analysis

S14: Data set: Calculation of relative fitness $\nu$

### SI Text

**S1.1: Effect of color markings on survival**

**Statistical analysis**

To test whether the color markings influenced the survival probability across treatments, we applied a univariate generalized mixed model (Ant nest ID and Group ID as random effects) using a binomial distribution and a logit link function.

**Results**

We could not find any effect of color on survival probability (model S18: $LR\chi^2_1 = 0.575$, estimate = 0.315, error = 0.415, p-value = 0.448, reference level: blue color, Fig. S1) across treatments.

**S1.2: Methods DNA extraction & preparation for sequencing (detailed version)**

1. DNA extraction: DNA was extracted using the DNeasy Blood and Tissue Kits from Qiagen following the manufacturer's protocol with minor modifications: Samples were lysed overnight but at least for six hours. AL and AE buffer was preheated until hand-warm before using them in step 2 and 8, respectively. At step 8, DNA was eluted twice with between 50 µl (for males) and 100 µl (for females) of AE buffer and then incubated for 10 to 15 minutes at room temperature in each time. Exact DNA concentrations were quantified using a Qubit fluorometer following the manufacturer's protocol. The remaining steps are a variation of the protocol as described in (1).

2. Sample dilution: All samples were normalized to a similar concentration of around 20 ng/µl by adding AE buffer. RE double digestion: 100 ng DNA (20 ng/µl) was double digested using two enzymes, MspI and MluCI, in equal concentrations (2 U) at 37 °C.

3. Adapter hybridization: Two types of single-stranded oligos were hybridized to double-stranded adapters by incubating them at 98 °C for 2.5 minutes and thereafter decreasing the temperature by 1 °C per minute. They contain an overhanging enzyme cut site-sequence, barcode, and a priming site for index primers. A total of seven different barcodes were used to account for the fact that 661 samples need to be distributed across seven 96-well plates.

4. Adapter ligation: The resulting double-stranded adapters were then ligated to the sticky end resulting from the enzyme digest using a different adapter for each of the seven 96-well plates. To this end, 100 ng of DNA of each sample was prepared together with 1.175 µM of the MspI adapter, 1.175 µM of the MluCI adapter and 100 U of T4 ligase and incubated at 23 °C for one hour and at 65 °C for ten minutes.

5. Test PCR: To check if both the enzyme digestion and adapter ligation was successful, samples were amplified and then ran through an electrophoresis gel. For the PCR 3 µl of the ligated DNA, 400 µM dNTPs, 0.25 µM of both For- and Rev-Primer, 0.5 U of OneTaq, 2 µl of 1x OneTag buffer and 4 µl distilled water were mixed for each sample. The PCR ran once for 30 seconds at 94 °C, then cycled 20 times through 15 seconds at 94 °C, 30 seconds at 55 °C and 45 seconds at 68 °C, then ended with 2 minutes at 68 °C. Based on the results of the electrophoresis, we had to exclude a total of 14 samples.

6. First sample pooling: The remaining samples were pooled from their original 96-well plate into one.

7. Sample clean-up: Samples were then cleaned up using an AMPure substitute. The DNA was mixed with twice the amount of beads and incubated for 5 minutes at room temperature. The DNA/bead solution was then placed on a magnetic rack, and the resulting supernatant was discarded. The remaining beads were washed twice with 100 µl EtOH and then air dried. Then, 16 µl $H_2O$ is added off the magnetic rack and incubated for another 5 minutes at room temperature before removing the clear supernatant which contained the DNA.

8. Index-PCR: Index primers with unique dual indices (UDI) were ligated to the DNA strands. To this end, 7.5 µl of the pooled and cleaned DNA, 12.5 µl of Q5/Phusion MM (2x), 5 µl of the index primers were combined. This was repeated for each well with DNA. The PCR ran once for 30 seconds at 98 °C, then cycled 8-12 times through 10 seconds at 98 °C, 75 seconds at 65 °C and then ended with 5 minutes at 65 °C.

9. Second sample pooling and size selection: 5 µl of each well were combined into a single 1.5 ml tube. For the size selection, a 1.5 % gel using Ultrapure agarose and TAE buffer was prepared. The ladder consisted of 10 µl of dye, 10 µl of a 100 kb DNA ladder and 40 µl of water. 90 µl of pooled DNA was mixed with 18 µl of dye. The gel ran for 2 hours at 90 V and thereafter the region between 320 and 500 kb was excised and dissolved using 4x Gel Dissolving Buffer at 37 °C. The DNA was then extracted by spinning it at

13000 g, then washing and spinning it twice with 200 µl of DNA Wash Buffer. Finally, after adding 20 µl of elution buffer, it was incubated for 1 min and then span for 1 min at 13000 g.

10. library quantification: We followed the manufacturer's protocol of the NEBNext Library Quant Kit.

##### S1.3 Effect of within-group kinship unweighted by sex ratio of the group

In addition to analyzing the mean kinship of females to their group members, of males to their group members, and among all group members which are weighted by the sex ratio of the group, we analyzed unweighted kinship levels among females, among males, and females and males (models S1-S17, Table S8) with results shown in S4-S7. All analyses were done analogous to the weighted kinship types. When we could not confirm Frequentist model assumptions, we additionally constructed Bayesian models.

### SI Figures

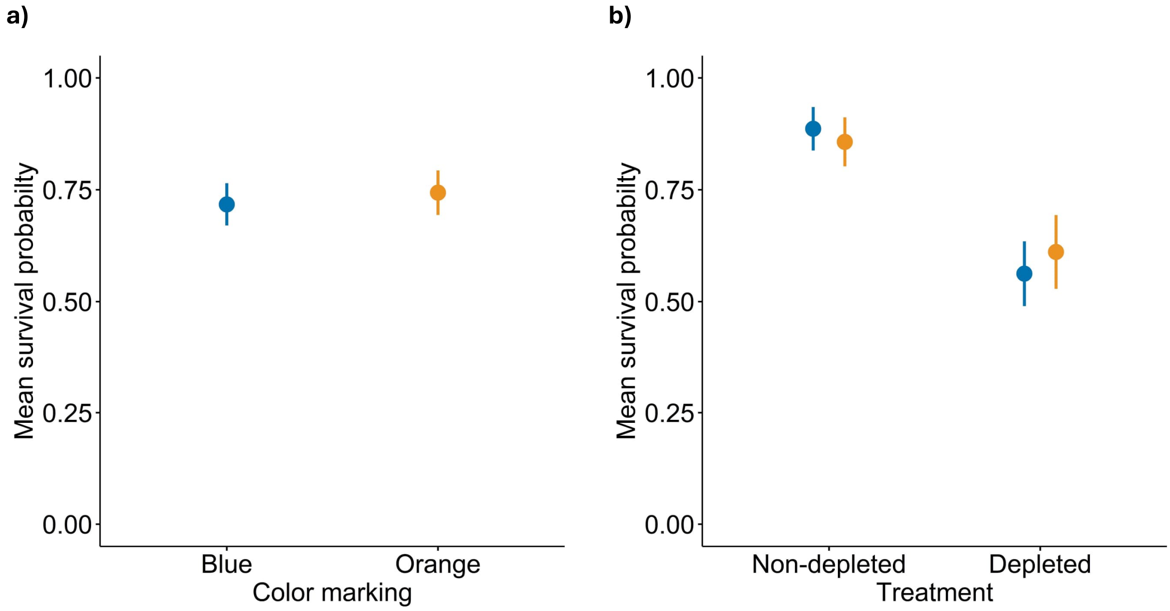


**Fig. S1.** Effects of color-marking on survival **a)** across treatment groups (blue: N = 92, orange: N = 78), **b)** and within treatment groups (blue/non-depleted: N = 44, blue/depleted: N = 48, orange/non-depleted: N = 42, orange/depleted: N = 36). Error bars indicate ±1 SEM.

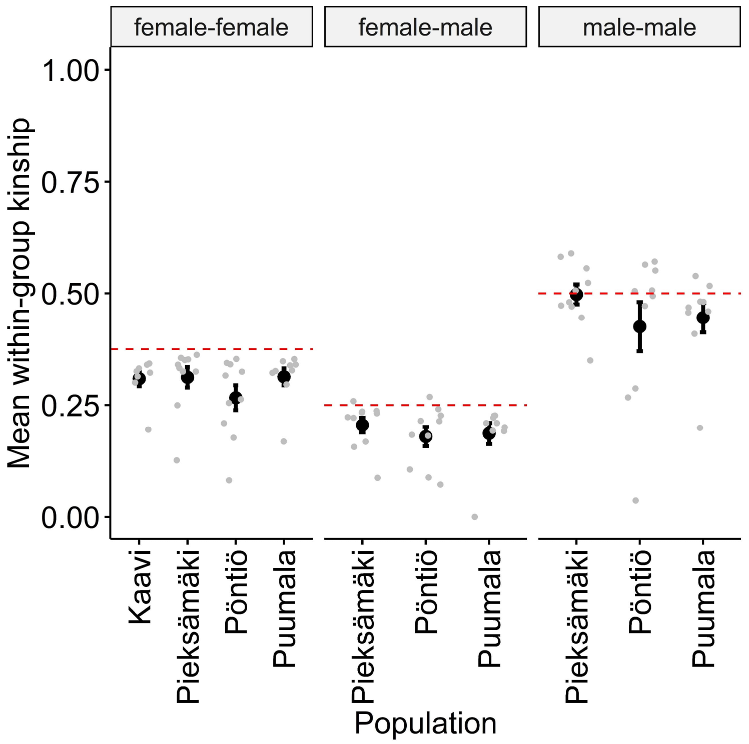


**Fig. S2.** Within-group kinships levels for female-female, female-male, and male-male comparisons of each population (in black) and larval group (in grey). Kaavi was excluded from the female-male and male-male comparisons due to low sample size of males (N = 1, and N = 2, respectively). Red lines indicate expected kinship value for full siblings. Error bars indicate ±1 SEM.

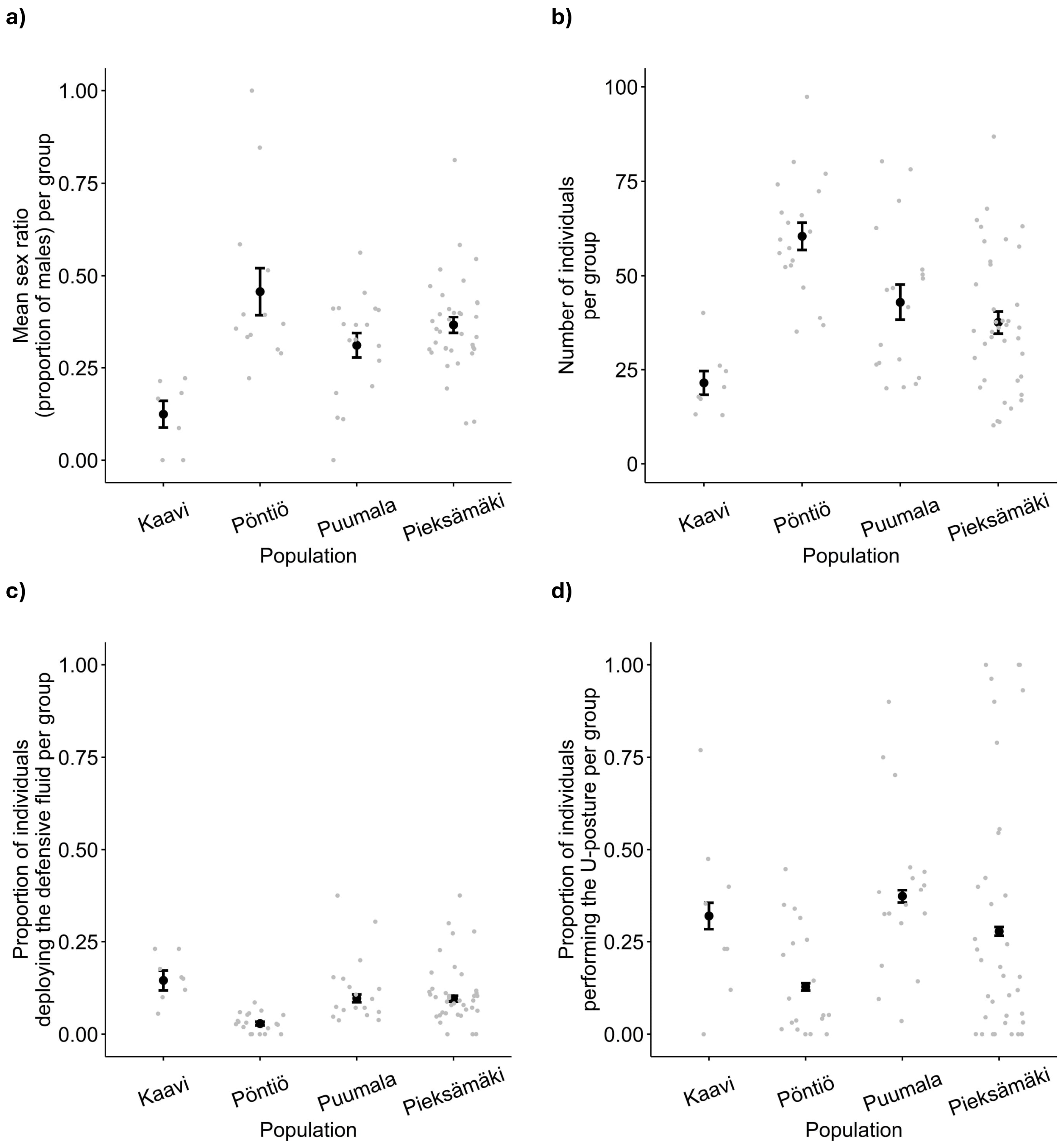


**Fig. S3.** Differences between populations in **a)** sex ratios, **b)** group size, **c)** deployment of the defensive fluid when an attack is simulated on an individual within the group, **d)** performing of the U-posture when an attack is simulated on an individual within the group. Each grey dot represents one larval group. Error bars indicate ±1 SEM.

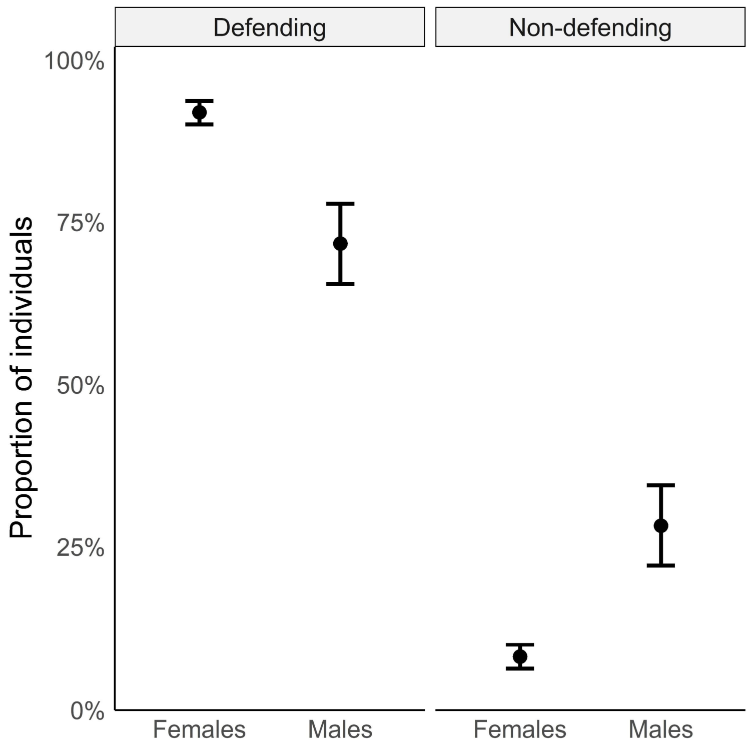


**Fig. S4.** Proportions of individuals deploying (defending) and not deploying (non-defending) the defensive fluid in each population when directly attacked (defending females: N = 204, defending males: N = 38, non-defending females: N = 18, non-defending males: N = 15). Error bars indicate ±1 SEM.

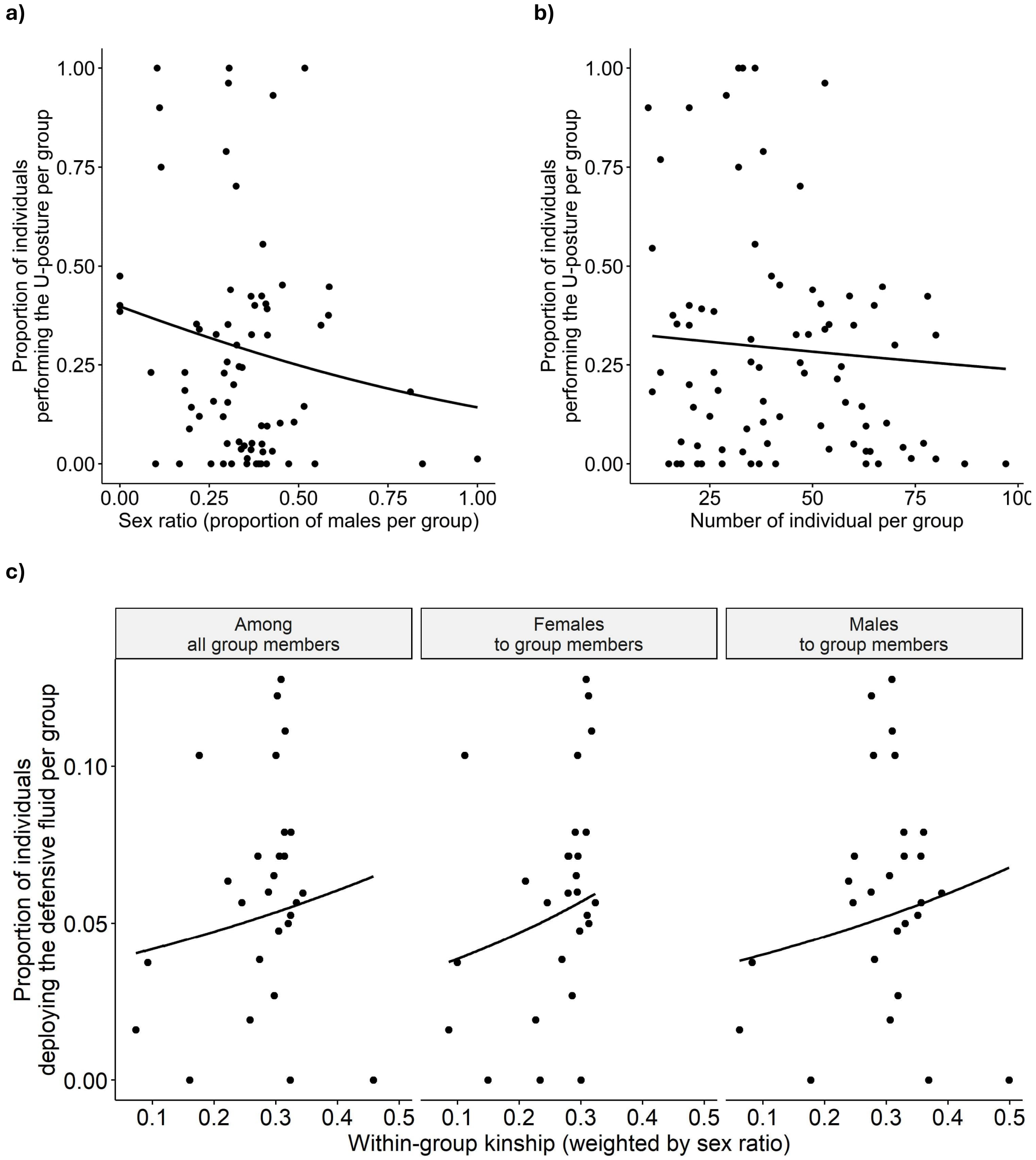


**Fig. S5.** Phenotypic association between participation in the defensive display (U-posture) when an attack is simulated on an individual within the group and the **a)** sex ratio (proportion of males) and **b)** size of the larval group. **c)** Phenotypic association between the contribution to the collective defense (secretion of the defensive fluid) when an attack is simulated on an individual within the group and the kinship level of the larval group.

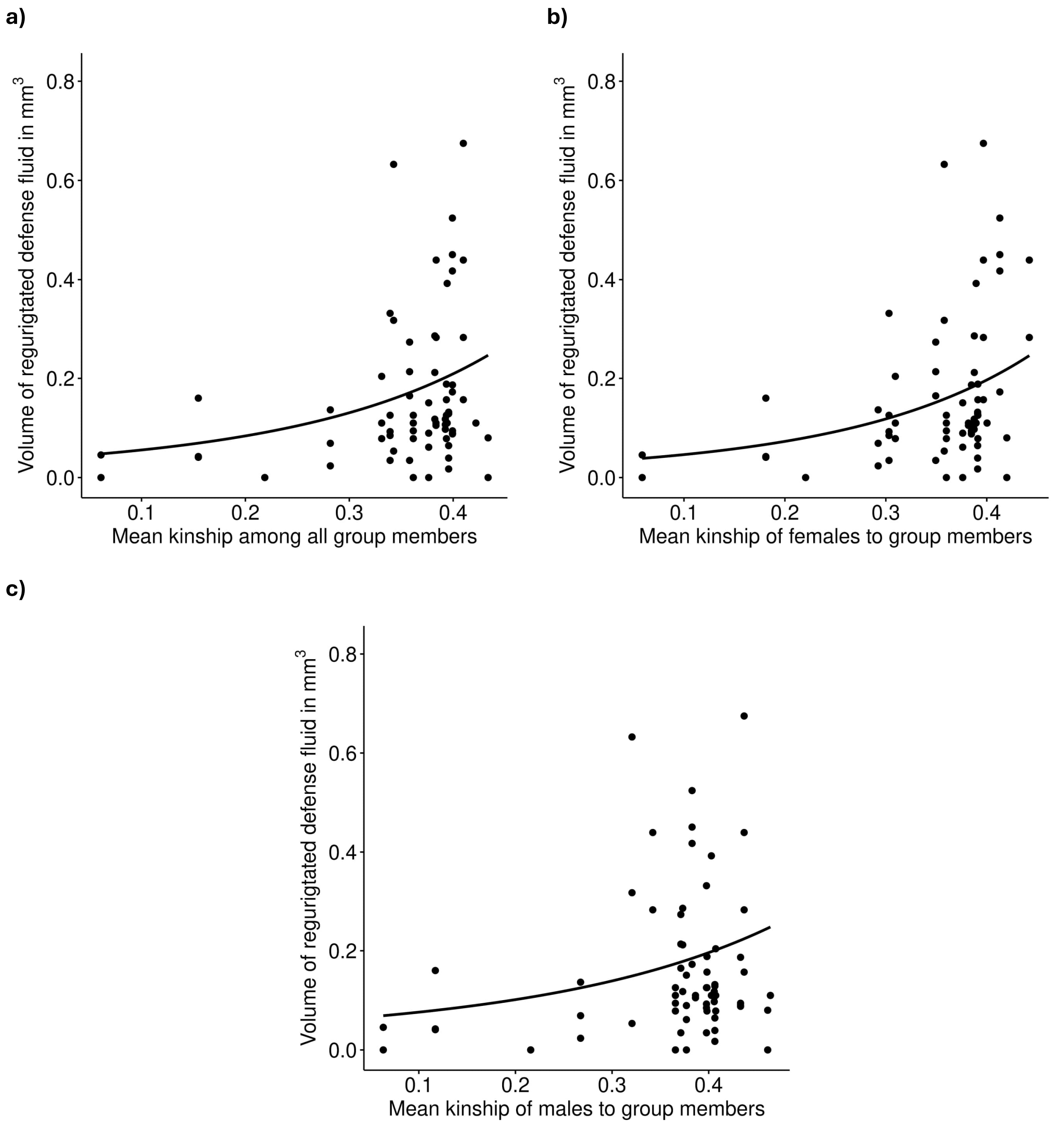


**Fig. S6.** Correlations between volume of deployed defensive fluid by females when directly attacked and **a)** mean kinship among all group members, **b)** mean kinship of females to group members, **c)** mean kinship of males to group members.

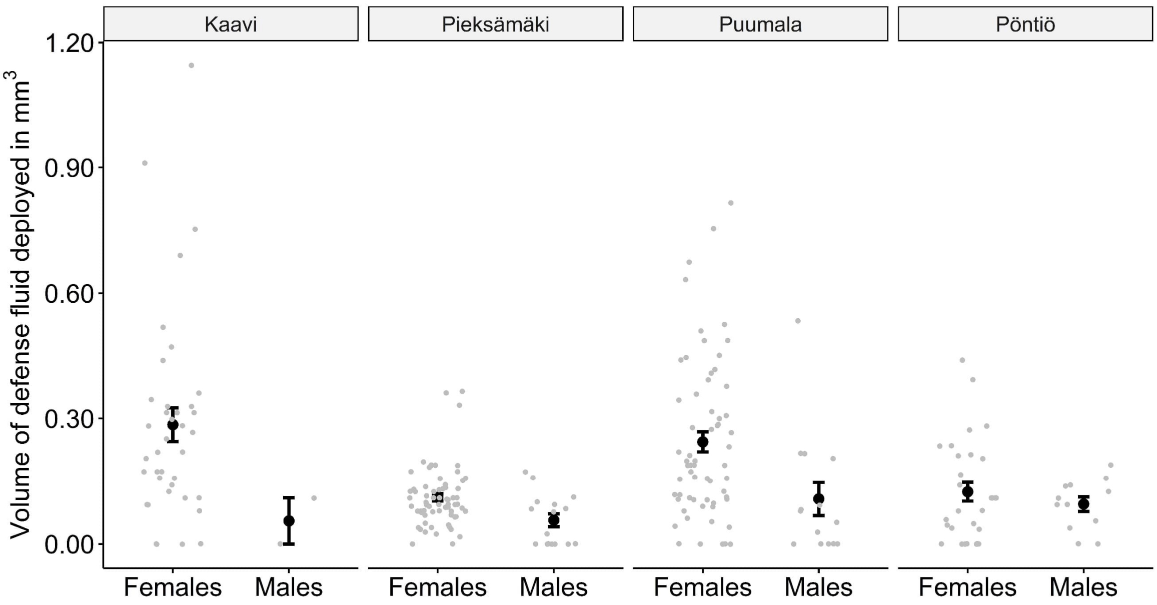


**Fig. S7.** Differences between populations and sexes in the volume of the deployed defensive fluid when directly attacked. Error bars indicate ±1 SEM. Each grey dot represents one individual.

a)

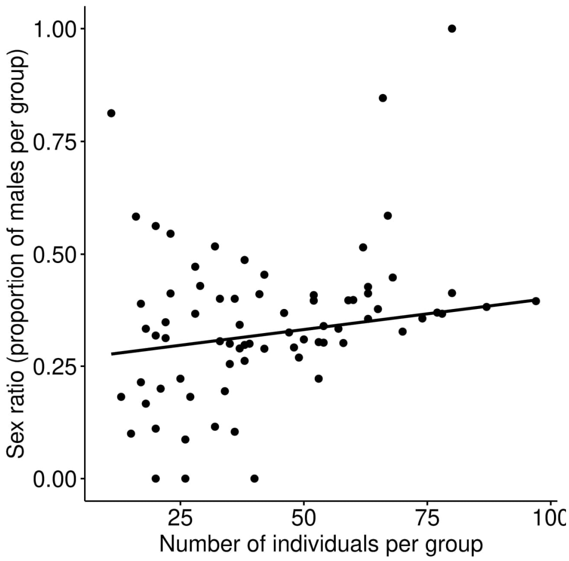


b)

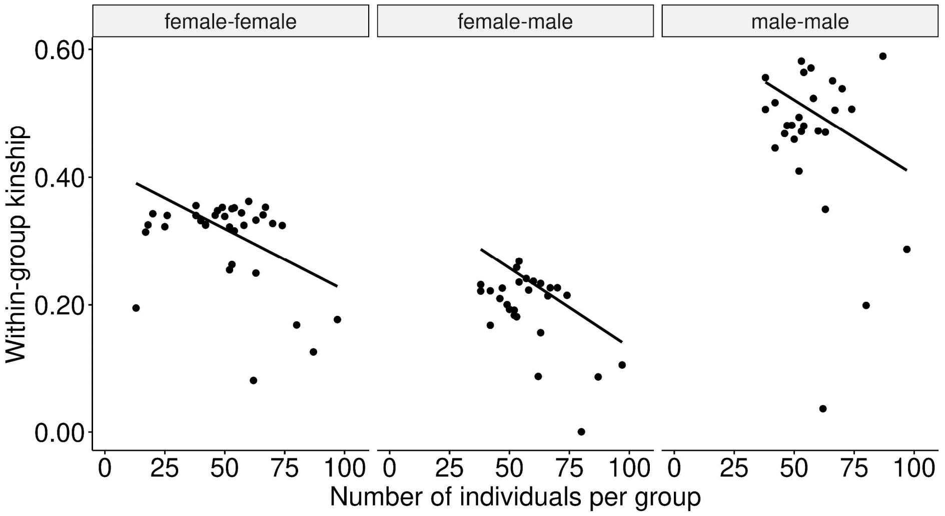


c)

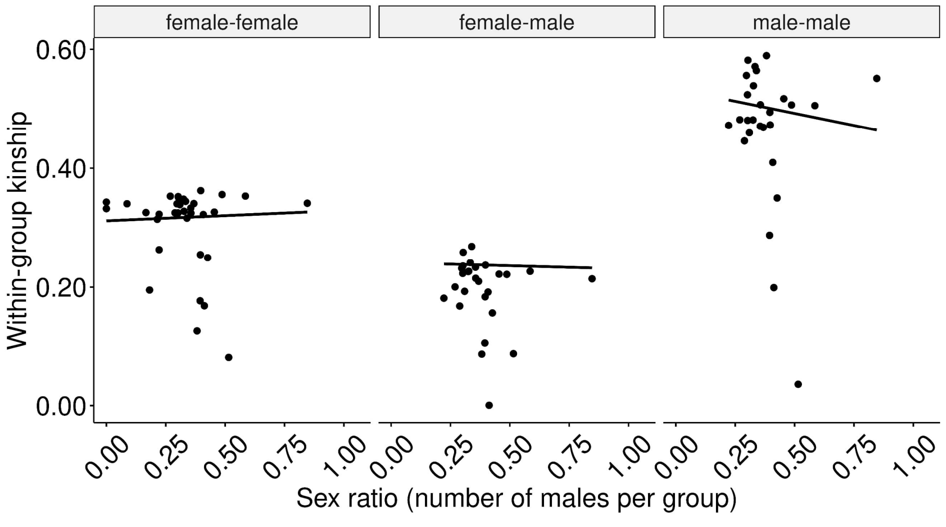


**Fig. S8.** Correlations between social environmental factors: **a)** group size and sex ratio, **b)** group size and within-group kinship, **c)** sex ratio and within-group kinship.

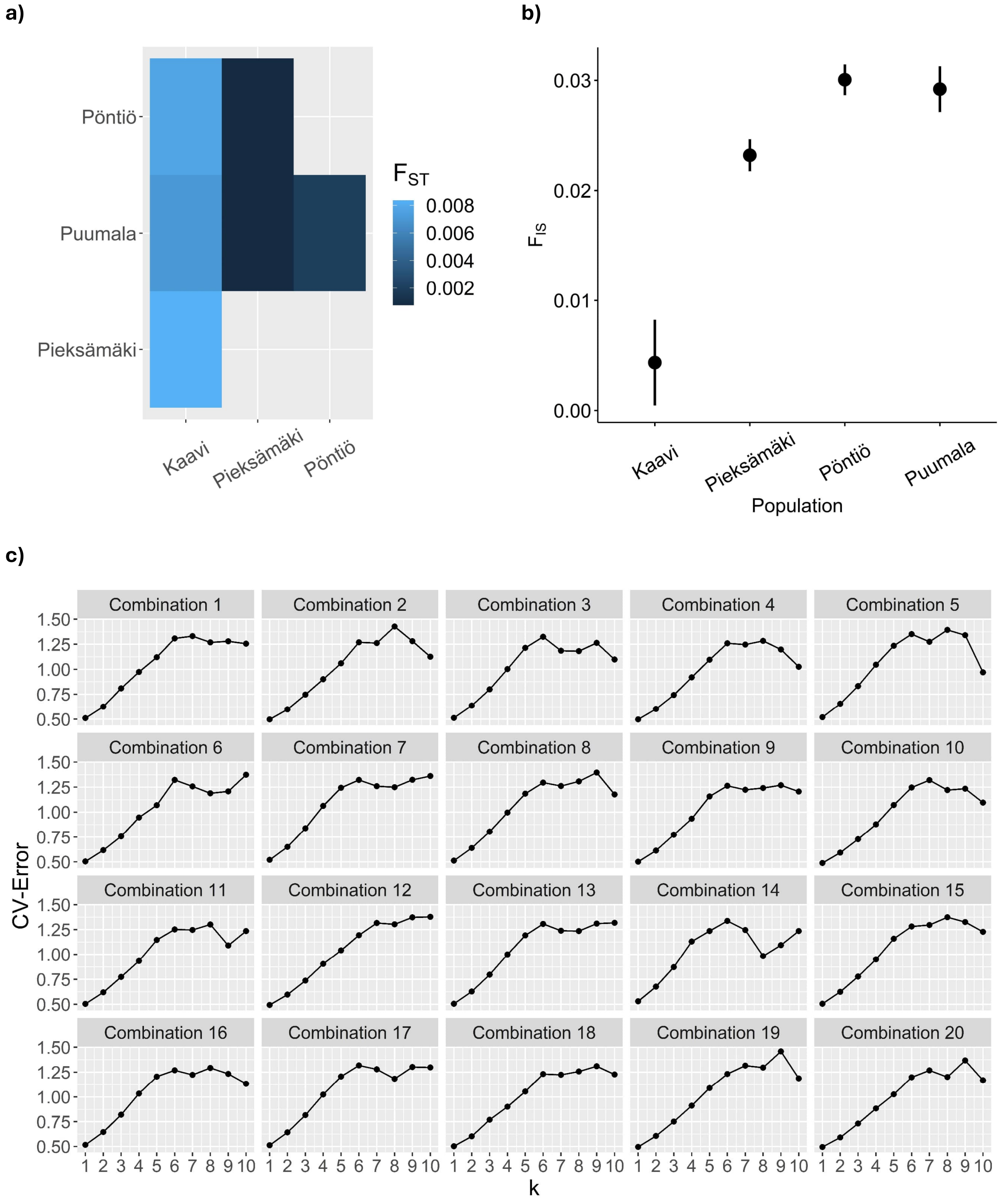


**Fig. S9. a)** Mean $F_{ST}$ values for all pairwise population comparisons. **b)** Mean $F_{IS}$ values for all pairwise population comparisons. Error bars indicate ±1 SEM **c)** CV-errors for different number of clusters/populations ranging from 1 to 10. All results are based on 20 random combinations of females (37 females per combination, 323 different females in total).

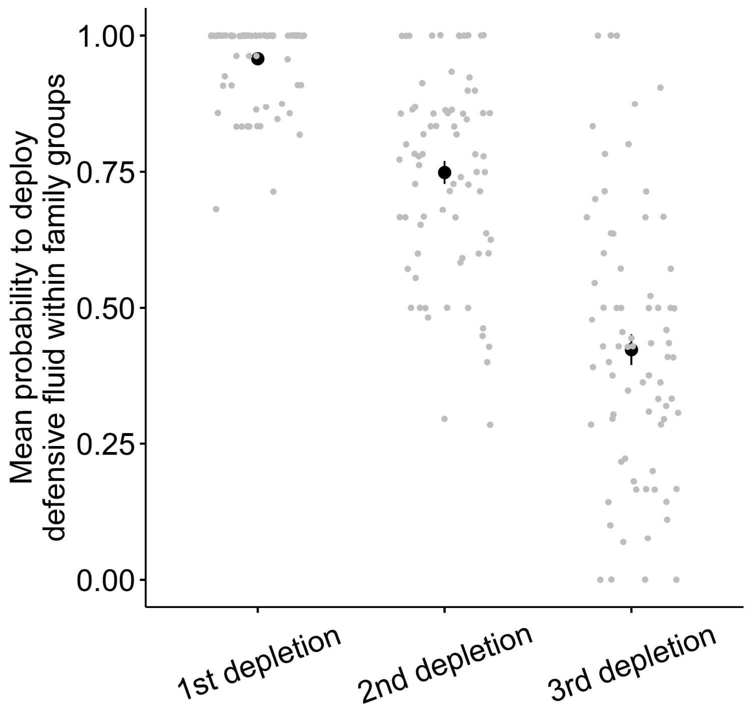


**Fig. S10.** Effect of the depletion treatment on the probability of the larvae to regurgitate the defensive fluid. These data are from 54 larval families (and 77 distinct larval groups in total) where individuals were depleted of their defensive fluid by poking each larva on their dorsal side and removing any regurgitated defensive fluid once a day for three days in a row. We recorded after each time whether they deployed any fluid or not (1st depletion: N = 1150, 2nd depletion: N = 1128, 3rd depletion: N = 1056). Error bars indicate ±1 SEM. Each grey dots represents one larval group.

#### SI Tables

**Table S1.** Differences between populations in individual and group-level traits. Significant values are in bold ($p < 0.05$).

| | | | | p-values of post-hoc test (Tukey Honest Significant Differences) | | | | | |
|---|---|---|---|---|---|---|---|---|---|
| **Model #** | **Trait** | **Chi²** | **p-values** | **Kaavi-Pieksämäki** | **Kaavi-Pöntiö** | **Kaavi-Puumala** | **Pieksämäki-Pöntiö** | **Pieksämäki-Puumala** | **Pöntiö-Puumala** |
| 3 | Proportion of individuals deploying the defensive fluid | LR $\chi^2_3$ = 53.39 | **<0.001** | 0.183 | **<0.001** | 0.247 | **<0.001** | 1 | **<0.001** |
| 4 | Proportion of individuals displaying the U-posture | LR $\chi^2_3$ = 15.948 | **0.001** | 0.997 | 0.086 | 0.876 | **0.011** | 0.904 | **<0.001** |
| 9 | Mean kinship among all group members | LR $\chi^2_3$ = 0.202 | 0.904 | - | - | - | - | - | - |
| 10 | Mean kinship of females to group members | LR $\chi^2_2$ = 1.919 | 0.383 | - | - | - | - | - | - |
| 11 | Mean kinship of males to group members | LR $\chi^2_3$ = 3.198 | 0.362 | - | - | - | - | - | - |
| 12 | Sex ratio | LR $\chi^2_3$ = 24.918 | **<0.001** | **0.001** | **<0.001** | **0.029** | 0.233 | 0.569 | **0.038** |
| 13 | Group size | LR $\chi^2_3$ = 36.01 | **<0.001** | 0.084 | **<0.001** | **0.021** | **0.001** | 0.688 | 0.014 |
| 5 | Defended/non-defended females | LR $\chi^2_3$ = 5.278 | 0.15 | - | - | - | - | - | - |
| 6 | Defended/non-defended males | LR $\chi^2_2$ = 1.448 | 0.485 | - | - | - | - | - | - |
| 7 | Defense volume females | LR $\chi^2_3$ = 27.995 (pop.)<br>LR $\chi^2_1$ = 0.446 (length) | **<0.001**<br>0.504 | **<0.001** | **0.004** | 0.649 | 0.991 | **0.010** | 0.012 |
| 8 | Defense volume males | LR $\chi^2_2$ = 1.368 (pop.)<br>LR $\chi^2_1$ = 5.306 (length) | 0.505<br>**0.021** | - | - | - | 0.763 | 0.968 | 0.471 |

Model 9 and 10 failed validations but results confirmed by Bayesian approach (Table S2).

**Table S2.** Re-run models with Bayesian approach: Differences between populations in kinship (weighted by sex ratio).

| | | Statistical values (estimate, error, low 95% CI, high 95% CI) | | |
|---|---|---|---|---|
| Model # | Trait | Pieksämäki-Pöntiö | Pieksämäki-Puumala | Pöntiö-Puumala |
| 9* | Mean kinship among all group members | 0.01 | -0.01 | 0.00 |
| | | 0.04 | 0.04 | 0.04 |
| | | -0.09 | -0.1 | -0.08 |
| | | 0.07 | 0.06 | 0.09 |
| 10* | Mean kinship of females to group members | -0.04 | -0.01 | -0.03 |
| | | 0.03 | 0.04 | 0.04 |
| | | -0.11 | -0.08 | -0.10 |
| | | 0.03 | 0.06 | 0.04 |

*Frequentist models (Table S1) failed validations. We therefore verified the results by additionally running Bayesian models.

**Table S3.** Correlations between sex ratio, group size, and kinship (unweighted by sex ratio). Variables whose 95% credible intervals do not include zero are in bold.

| Model # | Fixed factor | estimate | error | low 95% CI | high 95% CI | R-hat |
|---|---|---|---|---|---|---|
| Sex ratio | | | | | | |
| 40 | Group size | 0.0014 | -0.0011 | -0.0007 | 0.0035 | 1 |
| Female-female kinship | | | | | | |
| 41 | Sex ratio, | 0.0020 | 0.103 | -0.1860 | 0.2218 | 1 |
| | Group size | -0.0019 | 0.0009 | **-0.0037** | **-0.0002** | |
| Male-male kinship | | | | | | |
| 42 | Sex ratio, | -0.0082 | 0.221 | -0.517 | 0.3530 | 1 |
| | Group size | -0.0024 | 0.0019 | -0.0061 | 0.0013 | |
| Female-male kinship | | | | | | |
| 43 | Sex ratio, | -0.0096 | 0.0955 | -0.1970 | 0.1820 | 1 |
| | Group size | -0.0025 | 0.0008 | **-0.0041** | **-0.0009** | |

**Table S4.** Differences between populations in kinship (unweighted by sex ratio).

| Model # | Trait | $Chi^2$ | p-values |
|---|---|---|---|
| S1 | Female-female kinship | LR $\chi^2_3$ = 2.149 | 0.542 |
| S2 | Male-male kinship | LR $\chi^2_2$ = 0.992 | 0.609 |
| S3 | Female-male kinship | LR $\chi^2_3$ = 0.357 | 0.837 |

Models S1 and S2 failed validations but confirmed by Bayesian approach (Table S5).

**Table S5.** Re-run models with Bayesian approach: Differences between populations in kinship (unweighted by sex ratio).

| | | Statistical values (estimate, error, low 95% CI, high 95% CI) | | | | | |
|---|---|---|---|---|---|---|---|
| **Model #** | **Trait** | **Kaavi-Pieksämäki** | **Kaavi-Pöntiö** | **Kaavi-Puumala** | **Pieksämäki-Pöntiö** | **Pieksämäki-Puumala** | **Pöntiö-Puumala** |
| S1* | Female-female kinship | 0.00<br>0.04<br>-0.07<br>0.08 | -0.04<br>0.04<br>-0.11<br>0.04 | 0.01<br>0.04<br>-0.07<br>0.08 | 0.04<br>0.04<br>-0.03<br>0.11 | 0.00<br>0.04<br>-0.08<br>0.07 | -0.04<br>0.04<br>-0.12<br>0.03 |
| S3* | Female-male kinship | - | - | - | -0.01<br>0.03<br>-0.07<br>0.05 | -0.02<br>0.03<br>-0.08<br>0.04 | 0.01<br>0.03<br>-0.06<br>0.07 |

*Frequentist models (Table S3) failed validations. We therefore verified the results by additionally running Bayesian models.

**Table S6.** Effect of kinship (unweighted by sex ratio) on the proportion of individuals per group performing the U-posture or deploying the fluid. Significant values are in bold ($p < 0.05$).

| Model # | Fixed factor | Chi² | Beta | Error | p-value |
|---|---|---|---|---|---|
| Proportion of individuals displaying the U-posture within a group | | | | | |
| S4 | Female-female kinship | LR $\chi^2_1 = 3.184$ | 6.027 | 3.378 | 0.074 |
| S5 | Male-male kinship | LR $\chi^2_1 = 0.238$ | 1.008 | 2.067 | 0.626 |
| S6 | Female-male kinship | LR $\chi^2_1 = 2.410$ | 7.078 | 4.560 | 0.121 |
| Proportion of individuals deploying the defensive fluid within a group | | | | | |
| S7 | Female-female kinship | LR $\chi^2_1 = 0.872$ | 1.376 | 1.474 | 0.351 |
| S8 | Male-male kinship | LR $\chi^2_1 = 3.878$ | 2.373 | 1.205 | **0.048** |
| S9 | Female-male kinship | LR $\chi^2_1 = 0.977$ | 1.760 | 1.781 | 0.323 |

**Table S7.** Effect of kinship (unweighted by sex ratio) on individuals' defensive traits. Bold values indicate variables whose 95% credible intervals do not include zero.

| Model # | Fixed factor | Estimate | Error | Low 95% CI | High 95% CI | R-hat |
|---|---|---|---|---|---|---|
| Defensive fluid deployed when individually attacked | | | | | | |
| Females | | | | | | |
| S10 | Female-female kinship | 6.41 | 4.38 | -2.50 | 14.90 | 1 |
| S11 | Female-male kinship | 4.96 | 2.87 | -0.76 | 10.61 | 1 |
| Males | | | | | | |
| S12 | Male-male kinship | -6.90 | 11.51 | -32.94 | 12.44 | 1 |
| S13 | Female-male kinship | -0.65 | 6.73 | -15.75 | 11.21 | 1 |
| Volume of defensive fluid deployed when individually attacked | | | | | | |
| Females | | | | | | |
| S14 | Female-female kinship | 2.24 | 1.68 | -1.10 | 5.51 | 1 |
| | Length | 0.03 | 0.07 | -0.11 | 0.15 | |
| S15 | Female-male kinship | 3.65 | 1.09 | **1.38** | **5.70** | 1 |
| | Length | -0.04 | 0.08 | -0.19 | 0.11 | |
| Males | | | | | | |
| S16 | Male-male kinship | -3.21 | 6.72 | -17.66 | 8.53 | 1 |
| | Length | -0.22 | 0.20 | -0.62 | 0.19 | |
| S17 | Female-male kinship | -3.03 | 5.29 | -15.00 | 5.89 | 1 |
| | Length | -0.26 | 0.21 | -0.69 | 0.17 | |

**Table S8.** All models with model parameters.

| Model # | Model type | Response | Fixed factor | Random factor | Family | Link | iter warmup thin delta max_treedepth backend | Priors |
|---|---|---|---|---|---|---|---|---|
| 1 | glmmTMB | Survival | Treatment proportion | Ant nest ID, Group ID | Binomial | Logit | Not applicable | Not applicable |
| 2 | glmmTMB | Survival | Treatment proportion * treatment depleted | Ant nest ID, Group ID | Binomial | Logit | Not applicable | Not applicable |
| 3 | glmmTMB | Proportion of individuals deploying the fluid | Pop. ID | - | Binomial | Logit | Not applicable | Not applicable |
| 4 | glmmTMB | Proportion of individuals displaying the U-posture | Pop. ID | - | Beta-binomial | Logit | Not applicable | Not applicable |

*Table continues.*

| Model # | Model type | Response | Fixed factor | Random factor | Family | Link | iter<br>warmup<br>thin<br>delta<br>max_treedepth<br>backend | Priors |
|---|---|---|---|---|---|---|---|---|
| 5 | glmmTMB | Individual defense females (defended/ not defended) | Pop. ID | Group ID | Binomial | Logit | Not applicable | Not applicable |
| 6[1] | glmmTMB | Individual defense males (defended/ not defended) | Pop. ID | Group ID | Binomial | Logit | Not applicable | Not applicable |
| 7 | glmmTMB | Individual defense females (defensive fluid volume) | Pop. ID, Larval length | Group ID | Tweedie | Log | Not applicable | Not applicable |
| 8[1] | glmmTMB | Individual defense males (defensive fluid volume) | Pop. ID, Larval length | Group ID | Tweedie | Log | Not applicable | Not applicable |
| 9[1] | glmmTMB | Mean kinship among all group members | Pop. ID | Group ID | Gaussian | Identity | Not applicable | Not applicable |
| 9*[1] | glmmTMB | Mean kinship among all group members | Pop. ID | Group ID | Gaussian | Identity | 10000<br>5000<br>1<br>0.999<br>default<br>cmdstanr | Default |
| 10[1] | glmmTMB | Mean kinship of females to group members | Pop. ID | Group ID | Gaussian | Identity | Not applicable | Not applicable |
| 10*[1] | glmmTMB | Mean kinship of females to group members | Pop. ID | Group ID | Gaussian | Identity | 10000<br>5000<br>1<br>0.999<br>default<br>cmdstanr | Default |
| 11[1] | glmmTMB | Mean kinship of males to group members | Pop. ID | Group ID | Gaussian | Identity | Not applicable | Not applicable |
| 12 | glmmTMB | Sex ratio | Pop. ID | - | Gaussian | Identity | Not applicable | Not applicable |
| 13 | glmmTMB | Group size | Pop. ID | - | Gaussian | Identity | Not applicable | Not applicable |

*Table continues.*

| Model # | Model type | Response | Fixed factor | Random factor | Family | Link | iter<br>warmup<br>thin<br>delta<br>max_treedepth<br>backend | Priors |
|---|---|---|---|---|---|---|---|---|
| 14[1] | brm | Individual defense (defended/ not defended) | Sex | Pop. ID: Group ID | Binomial | Logit | 10000<br>5000<br>1<br>0.99<br>default<br>cmdstanr | Default |
| 15[1] | brm | Individual defense (defense volume) | Sex | Pop. ID: Group ID | Gamma | Log | 10000<br>5000<br>1<br>0.99<br>default<br>cmdstanr | Default |
| 16 | glmmTMB | Proportion of individuals displaying the U-posture | Sex ratio, Group size | Pop. ID | Beta-binomial | Logit | Not applicable | Not applicable |
| 17[1] | glmmTMB | Proportion of individuals displaying the U-posture | Mean kinship among all group members | Pop. ID | Beta-binomial | Logit | Not applicable | Not applicable |
| 18[1] | glmmTMB | Proportion of individuals displaying the U-posture | Mean kinship of females to group members | Pop. ID | Beta-binomial | Logit | Not applicable | Not applicable |
| 19[1] | glmmTMB | Proportion of individuals displaying the U-posture | Mean kinship of males to group members | Pop. ID | Beta-binomial | Logit | Not applicable | Not applicable |
| 20 | glmmTMB | Proportion of individuals deploying the fluid | Sex ratio, Group size | Pop. ID | Binomial | Logit | Not applicable | Not applicable |
| 21[1] | glmmTMB | Proportion of individuals deploying the fluid | Mean kinship among all group members | Pop. ID | Binomial | Logit | Not applicable | Not applicable |
| 22[1] | glmmTMB | Proportion of individuals deploying the fluid | Mean kinship of females to group members | Pop. ID | Binomial | Logit | Not applicable | Not applicable |

*Table continues.*

| Model # | Model type | Response | Fixed factor | Random factor | Family | Link | iter<br>warmup<br>thin<br>delta<br>max_treedepth<br>backend | Priors |
|---|---|---|---|---|---|---|---|---|
| 23[1] | glmmTMB | Proportion of individuals deploying the fluid | Mean kinship of males to group members | Pop. ID | Binomial | Logit | Not applicable | Not applicable |
| 24 | brm | Individual defense females (defended/ not defended) | Sex ratio, Group size | Pop. ID: Group ID | Binomial | Logit | 10000<br>5000<br>1<br>0.999<br>default<br>cmdstanr | Default |
| 25[1] | brm | Individual defense females (defended/ not defended) | Mean kinship among all group members | Pop. ID: Group ID | Binomial | Logit | 10000<br>5000<br>1<br>0.999<br>default<br>cmdstanr | Default |
| 26[1] | brm | Individual defense females (defended/ not defended) | Mean kinship of females to group members | Pop. ID: Group ID | Binomial | Logit | 10000<br>5000<br>1<br>0.999<br>default<br>cmdstanr | Default |
| 27[1] | brm | Individual defense females (defended/ not defended) | Mean kinship of males to group members | Pop. ID: Group ID | Binomial | Logit | 10000<br>5000<br>1<br>0.999<br>default<br>cmdstanr | Default |
| 28[1] | brm | Individual defense males (defended/ not defended) | Sex ratio, Group size | Pop. ID: Group ID | Binomial | Logit | 10000<br>5000<br>1<br>0.999<br>default<br>cmdstanr | Default |
| 29[1] | brm | Individual defense males (defended/ not defended) | Mean kinship among all group members | Pop. ID: Group ID | Binomial | Logit | 10000<br>5000<br>1<br>0.9999<br>default<br>cmdstanr | Default |
| 30[1] | brm | Individual defense males (defended/ not defended) | Mean kinship of females to group members | Pop. ID: Group ID | Binomial | Logit | 10000<br>5000<br>1<br>0.999<br>default<br>cmdstanr | Default |

*Table continues.*

| Model # | Model type | Response | Fixed factor | Random factor | Family | Link | iter<br>warmup<br>thin<br>delta<br>max_treedepth<br>backend | Priors |
|---|---|---|---|---|---|---|---|---|
| 31[1] | brm | Individual defense males (defended/ not defended) | Mean kinship of males to group members | Pop. ID: Group ID | Binomial | Logit | 10000<br>5000<br>1<br>0.999<br>default<br>cmdstanr | Default |
| 32 | brm | Individual defense females (defensive fluid volume + 0.001 | Sex ratio, Group size, Larval length | Pop. ID: Group ID | Gamma | Log | 10000<br>5000<br>1<br>0.99<br>default<br>cmdstanr | Default |
| 33[1] | brm | Individual defense females (defensive fluid volume + 0.001 | Mean kinship among all group members, Larval length | Pop. ID: Group ID | Gamma | Log | 10000<br>5000<br>1<br>0.999<br>default<br>cmdstanr | Default |
| 34[1] | brm | Individual defense females (defensive fluid volume + 0.001 | Mean kinship of females to group members, Larval length | Pop. ID: Group ID | Gamma | Log | 10000<br>5000<br>1<br>0.999<br>default<br>cmdstanr | Default |
| 35[1] | brm | Individual defense females (defensive fluid volume + 0.001 | Mean kinship of males to group members, Larval length | Pop. ID: Group ID | Gamma | Log | 10000<br>5000<br>1<br>0.999<br>default<br>cmdstanr | Default |
| 36[1] | brm | Individual defense males (defensive fluid volume + 0.001) | Sex ratio, Group size, Larval length | Pop. ID: Group ID | Gamma | Log | 10000<br>5000<br>1<br>0.9999<br>default<br>cmdstanr | Default |
| 37[1] | brm | Individual defense males (defensive fluid volume + 0.001) | Mean kinship among all group members | Pop. ID: Group ID | Gamma | Log | 10000<br>5000<br>1<br>0.999<br>default<br>cmdstanr | Default |

*Table continues.*

| Model # | Model type | Response | Fixed factor | Random factor | Family | Link | iter<br>warmup<br>thin<br>delta<br>max_treedepth<br>backend | Priors |
|---|---|---|---|---|---|---|---|---|
| 38[1] | brm | Individual defense males (defensive fluid volume + 0.001) | Mean kinship of females to group members | Pop. ID: Group ID | Gamma | Log | 10000<br>5000<br>1<br>0.999<br>default<br>cmdstanr | Default |
| 39[1] | brm | Individual defense males (defensive fluid volume + 0.001) | Mean kinship of males to group members | Pop. ID: Group ID | Gamma | Log | 10000<br>5000<br>1<br>0.999<br>default<br>cmdstanr | Default |
| 40 | brm | Group size | Sex ratio | Pop. ID | Gaussian | Identity | 10000<br>5000<br>1<br>0.999<br>15<br>cmdstanr | Default |
| 41 | brm | Female-female kinship | Group size, Sex ratio | Pop. ID | Gaussian | Identity | 10000<br>5000<br>1<br>0.9999<br>15<br>cmdstanr | Default |
| 42[2] | brm | Male-male kinship | Group size, Sex ratio | Pop. ID | Gaussian | Identity | 10000<br>5000<br>1<br>0.9999<br>15<br>cmdstanr | Default |
| 43[2] | brm | Female-male kinship | Group size, Sex ratio | Pop. ID | Gaussian | Identity | 10000<br>5000<br>1<br>0.9999<br>15<br>cmdstanr | Default |
| S1 | glmmTMB | Female-female kinship | Pop. ID | - | Tweedie | Log | Not applicable | Not applicable |
| S1* | brm | Female-female kinship | Pop. ID | - | Gaussian | Identity | 10000<br>5000<br>1<br>0.99<br>default<br>cmdstanr | Default |
| S2[2] | glmmTMB | Male-male kinship | Pop. ID | - | Tweedie | Log | Not applicable | Not applicable |
| S3[2] | glmmTMB | Female-male kinship | Pop. ID | - | Tweedie | Log | Not applicable | Not applicable |

*Table continues.*

| Model # | Model type | Response | Fixed factor | Random factor | Family | Link | iter<br>warmup<br>thin<br>delta<br>max_treedepth<br>backend | Priors |
|---|---|---|---|---|---|---|---|---|
| S3*[2] | brm | Female-male kinship | Pop. ID | - | Gaussian | Identity | 10000<br>5000<br>1<br>0.99<br>default<br>cmdstanr | Default |
| S4 | glmmTMB | Proportion of individuals displaying the U-posture | Female-female kinship | Pop. ID | Beta-binomial | Logit | Not applicable | Not applicable |
| S5[2] | glmmTMB | Proportion of individuals displaying the U-posture | Male-male kinship | Pop. ID | Beta-binomial | Logit | Not applicable | Not applicable |
| S6[2] | glmmTMB | Proportion of individuals displaying the U-posture | Female-male kinship | Pop. ID | Beta-binomial | Logit | Not applicable | Not applicable |
| S7 | glmmTMB | Proportion of individuals deploying the fluid | Female-female kinship | Pop. ID | Binomial | Logit | Not applicable | Not applicable |
| S8[2] | glmmTMB | Proportion of individuals deploying the fluid | Male-male kinship | Pop. ID | Binomial | Logit | Not applicable | Not applicable |
| S9[2] | glmmTMB | Proportion of individuals deploying the fluid | Female-male kinship | Pop. ID | Binomial | Logit | Not applicable | Not applicable |
| S10 | brm | Individual defense females (defended/ not defended) | Female-female kinship | Pop. ID: Group ID | Binomial | Logit | 10000<br>5000<br>1<br>0.999<br>default<br>cmdstanr | Default |
| S11[2] | brm | Individual defense females (defended/ not defended) | Female-male kinship | Pop. ID: Group ID | Binomial | Logit | 10000<br>5000<br>1<br>0.999<br>default<br>cmdstanr | Default |
| S12[2] | brm | Individual defense males (defended/ not defended) | Male-male kinship | Pop. ID: Group ID | Binomial | Logit | 10000<br>5000<br>1<br>0.9999<br>default<br>cmdstanr | Default |
| S13[2] | brm | Individual defense males (defended/ not defended) | Female-male kinship | Pop. ID: Group ID | Binomial | Logit | 10000<br>5000<br>1<br>0.999<br>default<br>cmdstanr | default |

*Table continues.*

| Model # | Model type | Response | Fixed factor | Random factor | Family | Link | iter<br>warmup<br>thin<br>delta<br>max_treedepth<br>backend | Priors |
|---|---|---|---|---|---|---|---|---|
| S14 | brm | Individual defense females (defensive fluid volume + 0.001 | Female-female kinship, Larval length | Pop. ID: Group ID | Gamma | Log | 10000<br>5000<br>1<br>0.999<br>default<br>cmdstanr | default |
| S15[2] | brm | Individual defense females (defensive fluid volume + 0.001 | Female-male kinship, Larval length | Pop. ID: Group ID | Gamma | Log | 10000<br>5000<br>1<br>0.999<br>default<br>cmdstanr | default |
| S16[2] | brm | Individual defense males (defensive fluid volume + 0.001) | Male-male kinship, Larval length | Pop. ID: Group ID | Gamma | Log | 10000<br>5000<br>1<br>0.999<br>default<br>cmdstanr | default |
| S17[2] | brm | Individual defense males (defensive fluid volume + 0.001) | Female-male kinship, Larval length | Pop. ID: Group ID | Gamma | Log | 10000<br>5000<br>1<br>0.999<br>default<br>cmdstanr | default |
| S18 | glmmTMB | Survival | Color | Ant nest ID, Group ID | Binomial | Logit | Not applicable | Not applicable |

*Frequentist model failed validations. We therefore verified the results by additionally running Bayesian models.
[1]Kaavi was excluded because of the very low sample size for males (N = 1 for mean kinship among all group members, N = 1 for mean kinship of males to other group members, and N = 2 for mean kinship of females to other group members, N = 2 for male individual-level traits).
[2]Kaavi was excluded because of the very low samples size for males (N = 2 for female-male comparisons and N = 1 for male-male comparisons, N = 2 for male individual-level traits)
